# Promoted Read-through and Mutation Against Pseudouridine-CMC by an Evolved Reverse Transcriptase

**DOI:** 10.1101/2024.07.03.601893

**Authors:** Zhiyong He, Weiqi Qiu, Huiqing Zhou

## Abstract

Pseudouridine (Ψ) is an abundant RNA chemical modification that can play critical roles in the biological functions of RNA, and RNA-therapeutic applications. Current Ψ detection methods are limited in identifying Ψs at base-resolution in U-rich sequence contexts, where Ψ occurs frequently. The N-cyclohexyl N’-(2-morpholinoethyl)carbodiimide (CMC) can selectively label Ψ in RNA by forming the CMC-Ψ adduct. Here we report that an evolved reverse transcriptase (“RT-1306”) shows promoted read-through and mutation against the CMC-Ψ. The mutation signature can resolve the occurrence of Ψs within UU-containing sequences. We developed “Mut-Ψ-seq” utilizing CMC and RT-1306 for transcriptome-wide mapping of Ψ at base-resolution. The mutation signatures robustly identify reported Ψs in human rRNAs via the ROC analysis, and elongated CMC reaction duration increases the detection sensitivity of Ψ. We report a high-confidence list of Ψ sites in polyA-enriched RNAs from HEK-293T cells identified by orthogonal chemical treatments (CMC and bisulfite). The mutation signatures resolve the position of Ψ in UU-containing sequences, revealing diverse occurrence of Ψs in such sequences. This work provides new methods and datasets for biological research of Ψ, and demonstrates the potential of combining the reverse transcriptase engineering and selective chemical labeling to expand the toolkit for RNA chemical modifications studies.

## INTRODUCTION

Pseudouridine (Ψ), an isomer of uridine, is the first chemically modified ribonucleotide identified in RNA noted as “the fifth nucleotide” in 1957,^[1]^ and an abundant RNA chemical modification occurring in all three domains of life.^[2]^ Endogenous Ψs are known to be installed by stand-alone or RNA-dependent pseudouridine synthases.^[3]^ To form a Ψ, a uridine (U) undergoes isomerization starting with the cleavage of the glycosidic bond (N1-C1’), followed by a 180° rotation of the base around the N3-C6 axis, and the reformation of a C5-C1’ linkage between the rotated uracil and the ribose (**Figure 1a**). Ψ presents the same Watson-Crick-Franklin base-pairing face as U, and contains an extra H-bond donor (N1-H1) and a C-C atypical glycosidic linkage. The chemical structure of Ψ endows unique features in modulating base stacking energetics,^[4]^ stability,^[5]^ conformation,^[6]^ and molecular recognition^[7]^ of Ψ-modified RNAs. While Ψ was initially known to occur in non-coding RNAs such as rRNAs, tRNAs, and snRNAs, growing evidence over the last decade revealed that Ψ is an abundant modification occurring in mRNAs and long non-coding RNAs that play regulatory functions in gene expression and diseases.^[8]^ The occurrence of endogenous Ψs can alter in response to external stress conditions suggesting potential dynamics in Ψ regulation.^[8b, 8c, 9]^

**Figure 1.**
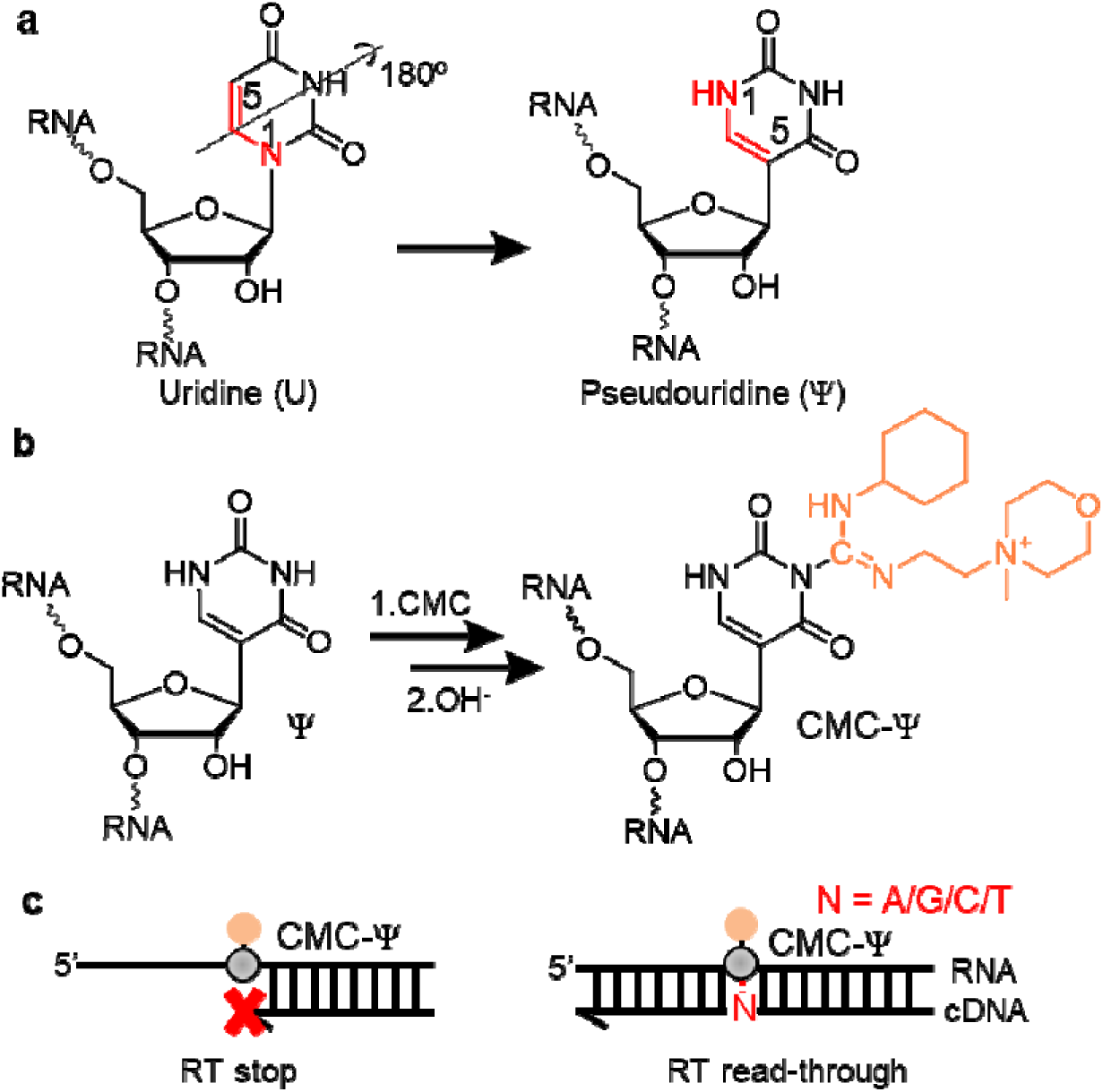
Structure and CMC-based detection of Ψ in RNA. **a**) Chemical structures showing the formation of Ψ upon the isomerization of U. **b**) Reaction scheme of CMC with Ψ yielding the N3 CMC-Ψ adduct (“CMC-Ψ”). **c**) Illustrations of RT stop and read-through events against the CMC-Ψ.

Accurate and precise detection of the occurrence of Ψs in the transcriptome is crucial for identifying the functional context of Ψ modification.^[10]^ There have been rapid advances in the transcriptome-wide mapping methods of RNA chemical modifications using RNA-seq based technologies.^[11]^ In order to map Ψs, “Pseudo-seq”^[8a]^ and “Ψ-seq”^[8b]^ coupled the classic “CMC” reaction^[12]^ and RNA-seq to map Ψ at single-base resolution. Briefly, biological RNAs were treated with N-cyclohexyl N’-(2-morpholinoethyl)carbodiimide (CMC), followed by an alkaline treatment step.^[12]^ CMC reacts with deprotonated Ψ-N1, Ψ-N3, U-N3, and G-N1 and forms corresponding CMC-adducts. The adducts Ψ-N1-CMC, U-N3-CMC, and G-N1-CMC get readily reverted back to U-N3-H3, G-N1-H1, and Ψ-N1-H1 under alkaline condition; the Ψ-N3-CMC is resistant to alkaline hydrolysis likely due to the presence of the negative charge at N1 (**Figure 1b**).^[8a, 8b, 12]^ The resulted Ψ-N3-CMC presents a bulky and positively charged group at the canonical base-pairing interface and thus promotes stop signatures during reverse transcription (RT); RT stops were subsequently measured by next-generation sequencing and used as signatures to identify Ψ (**Figure 1c**).^[13]^ However, RT stop signatures were subjected to high background noise due to non-random RNA fragmentation, ligation biases, RNA degradation, and stably folded RNA structures. These factors yielded a high false positive rate and low data reproducibility for Ψ identification in low-abundance RNAs, such as mRNAs and long non-coding RNAs. Moreover, RT stop signatures often fail to identify consecutive or clustered Ψs. ^[8a, 14]^

Recently, “RBS-Seq”,^[15]^ “BID-seq”,^[16]^ and “PRAISE”^[17]^ were developed for Ψ mapping utilizing bisulfite/sulfite treatment to RNA, which produced bisulfite-Ψ adducts and thus RT deletion signatures at bisulfite-Ψs. Bisulfite-based methods demonstrated advanced capabilities in mapping multiple modifications and improved detection sensitivity and reproducibility of Ψ.^[15-17]^ Yet, one caveat for using RT deletion signature is that deletion cannot accurately determine the location of Ψ in any UU sequence context.^[16-17]^ Indeed, Ψ was frequently identified in UU-containing sequences in polyA-enriched RNAs from human cell lines. For instance, 1357 of the total 2209 Ψ sites (61%) reported by PRAISE occurred in UU-containing sequences in polyA-enriched RNAs from HEK-293T cells, identified within the UU-containing sequence range rather than at single-base resolution;^[17]^ BID-seq reported the identification of GUUC and poly-U (five Us or more) motifs for Ψ occurrence^[16]^ and a recently reported Nanopore-based Ψ detection method revealed that the majority of Ψ-containing sequence motifs contained UU sequence contexts.^[18]^

Despite the wide occurrence, the biosynthesis and function for Ψs in UU-containing sequences have yet been established. Several studies showed evidence for the installation of Ψ in the GUΨC motif in mRNAs by the human pseudouridine synthase TRUB1,^[14, 16-17]^ which was known to recognize and introduce Ψ55 in the GUΨC motif in the T-loop of tRNAs.^[19]^ The biosynthesis of Ψs occurring in other UU-containing sequences in polyA-enriched RNAs remain unclear. Knockdown studies show that these Ψs can be partially accounted by the RNA-guided dyskerin pseudouridine synthase 1 (DKC1); however, the guide RNA sequences have yet been reported and many Ψs sites have no identified writers.^[17]^ In addition to biosynthesis, Ψ’s occurrence in different codon positions within UU-containing codon sequences altered the translation error rate demonstrated by *in vitro* translation assay.^[20]^ Resolving the detection challenges of Ψ in U-rich sequence context is critical for understanding the biosynthetic mechanism and regulatory function in gene expression of pseudouridination in mRNAs.^[18]^

Here we report that a recently evolved reverse transcriptase (RTase) “RT-1306”^[21]^ shows promoted read-through efficiency and mutation rates when processing CMC-Ψ adduct in CMC-treated RNA (**Figure 1c**). We developed “Mut-Ψ-seq” utilizing RT-1306 in conjunction with the classic CMC reaction, to map Ψ at single-base resolution in the transcriptome. Mut-Ψ-seq data showed excellent performance in identifying the reported Ψ sites in rRNAs^[22]^ by mutation signatures of RT-1306 via the receiver operating characteristic (ROC) curve analysis. Ψ-identification can be affected by the CMC reaction condition. Sequencing results showed elongated CMC reaction duration (2-hour) promoted the RT signatures significantly at Ψ sites compared to the 20-minute reaction, which can improve the detection sensitivity for Ψ validation efforts. However, the 2-hour reaction duration raised concerns about increased RNA loss and elevated background signatures on unmodified Us, which didn’t out-perform the 20-minute CMC reaction in terms of Ψ-identification efficiency according to the ROC curve analysis. We then used ROC-guided criteria to identify Ψs in the transcriptome and reported a high-confidence list of 44 Ψ sites in abundant mRNAs and non-coding RNAs identified by orthogonal chemical treatments: CMC for Mut-Ψ-seq and the bisulfite for PRAISE.^[17]^ 77% of these sites occurred in UU-containing sequences, and the RT mutation signature resolved the position of Ψs in most UU sequence contexts. Mut-Ψ-seq has broad applications for transcriptome-wide mapping and locus-specific detection of Ψ in any sequence contexts, and for investigations of biosynthesis mechanisms and regulatory functions of Ψ.

## RESULTS AND DISCUSSION

### RT-1306 shows promoted read-through and mutation against CMC-Ψ in RNA oligonucleotides

To examine the read-through propensities of RT-1306 against the CMC-Ψ adduct, we first prepared RNA oligonucleotides (Ψ-oligo1 and Ψ-oligo2, sequences shown in **Table S1**) that carry a single Ψ in the sequence into the CMC-Ψ RNA through the reported CMC reaction condition.^[23]^ RNA oligos were treated with excess amount of CMC for 16 hours followed with the alkaline treatment step (**Figure 1b** and **Methods**). We performed direct electrophoresis analysis of RNA product after CMC reaction, where the RNA product showed as a smeared band, consistent with addition of CMC on the Ψ, Us and Gs. In contrast, the smeary feature disappeared after alkaline treatment, indicating the removal of the CMC group on Us and Gs (**Figure S1**).^[24]^

We then applied Superscript III (SSIII) RT stop assay to examine the products of the RT reaction with fluorescein amidite (FAM)-labeled primers (**Methods**). SSIII was reported to present a near-complete RT stop at the 100% CMC-Ψ site under regular RT condition (with Mg^2+^);^[23]^ our data showed ∼90% RT stop at the CMC-Ψ site quantified by fluorescence intensity of the gel bands, indicating near-complete Ψ into CMC-Ψ conversion in both CMC treated Ψ-oligo1 and Ψ-oligo2 (**Figure 2a** and **Figure S2**). We did not observe additional bands except for the residual primer, RT stop and full-length cDNAs, suggesting that there are no major remaining side products of CMC-U and CMC-G adducts on the RNA oligos after the base-treatment (**Figure 2a** and **Figure S2**).

**Figure 2.**
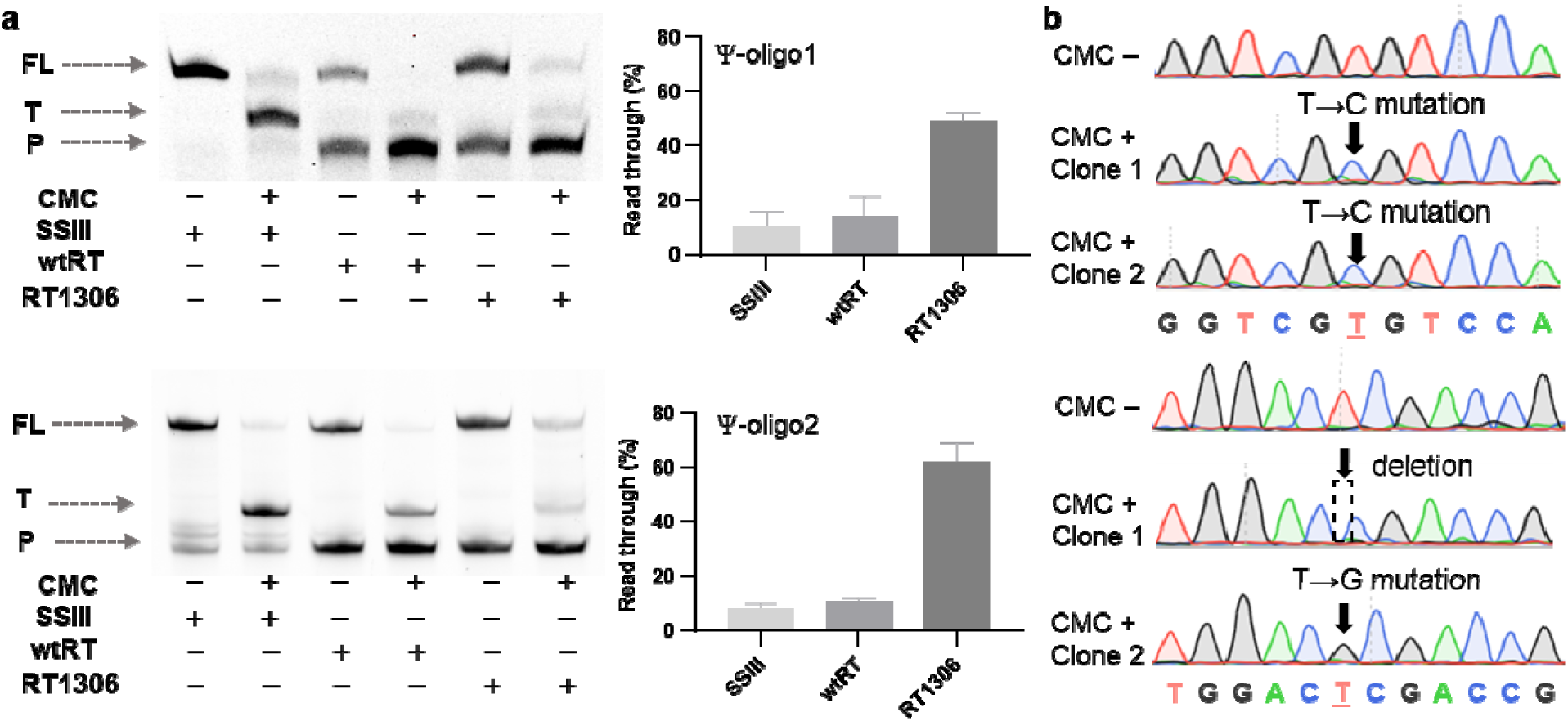
Promoted read-through efficiencies and mutations against CMC-Ψ by RT-1306. **a**) Shown on the left are the fluorescence images of RT stop assay gels for SSIII, wtRT, and RT-1306 reading against Ψ-oligo1 and Ψ-oligo2 RNAs with and without CMC treatment. RT products were run and separated on 15% 8M Urea-PAGE gels and imaged by the fluorescence imaging. Positions of the FAM-labeled RT primer, truncated cDNA at the Ψ site, and the full-length cDNA products are labeled with “P”, “T” and “FL”, respectively. The same gels were also stained by SYBR-Gold and imaged (**Figure S2**). Shown on the right are quantified RT read-through based on the RT stop assay. The RT read-through efficiency over CMC-Ψ were quantified by the ratio of the fluorescence intensity of the “T” band over the sum of intensities of the “T” and “F” bands. Error bars represent the standard deviation of n = 2 replicates; full gel images of 2 replicates are presented in **Figure S2**. **b**) Colony sequencing data of the cDNA products from RT1306 processing Ψ-oligo1 and Ψ-oligo2 RNAs with (“CMC+”) and without (“CMC-”) CMC treatments (**Methods**). Shown are the clones that carry mutations at the Ψ sites and data for all sequenced colonies are shown in **Figure S3**.

Next, we assessed the read-through propensity of RT-1306 against the two CMC treated Ψ oligonucleotides (CMC-Ψ-oligo1 and CMC-Ψ-oligo2, sequences shown in **Table S1**), via the RT stop assay with RT-1306. We used the p66 subunit of wild-type HIV-RT (wtRT) as a control. The wtRT showed ∼ 94% RT stop at the CMC-Ψ site, comparable to SSIII (**Figure 2a**). Interestingly, RT-1306 showed 46-53% read-through efficiencies over CMC-Ψ for both oligos, ∼4-fold higher than the those of wtRT and SSIII (**Figure 2a** and **Figure S2**). It was reported in the literature that the RT read-through efficiency against CMC-Ψ can be promoted by adding Mn^2+^;^[13]^ notably, RT-1306 showed improved read-through under the regular Mg^2+^-based RT conditions, demonstrating unique advances by RTase evolution.^[21]^ Despite that RT-1306 was engineered against m^1^A, it showed extended applications in to the CMC-Ψ adduct which carries a charged and more bulky modification.

To assess whether RT-1306 produced any signature that can be deployed to identify Ψ in a read-through cDNA product, we performed colony sequencing assay to characterize the sequence of the read-through RT product by RT-1306 (**Methods** and **Figure S3a**). 2 out of the total 10 sequenced colonies sequenced for CMC-Ψ-oligo1 showed T→C mutations. Among the total 7 sequenced colonies for CMC-Ψ-oligo2, one showed a T→G mutation and two showed deletions at the Ψ position (**Figure 2b**, **Figure S3b** and **Table S1**). This indicated that RT-1306 was capable of generating signatures to identify Ψ in RNA, which made it promising to be applied for CMC-based Ψ mapping. The colony sequencing assay provided a convenient method for site-directed detection of Ψs.^[13, 25]^

### Ψ into CMC-Ψ conversion and RNA loss by CMC treatment

Before applying the RT-1306 directly into Ψ-seq, we noticed that the 16-hour CMC treatment condition^[23]^ resulted in significant RNA loss; only 2% of the input RNA oligonucleotides were recovered after the CMC reaction and the alkaline treatment step (**Figure 3**). Our data suggested the loss of RNA already occurred at the first CMC reaction step (**Figure S4a**). Indeed, the reported CMC-based Ψ-sequencing methods applied considerably shorter reaction time for the CMC reaction step: 20-30 minutes (**Figure S4b**).^[8a-c]^ However, neither the conversion rate of Ψ into CMC-Ψ, nor the level of RNA loss, were reported under these conditions.

**Figure 3.**
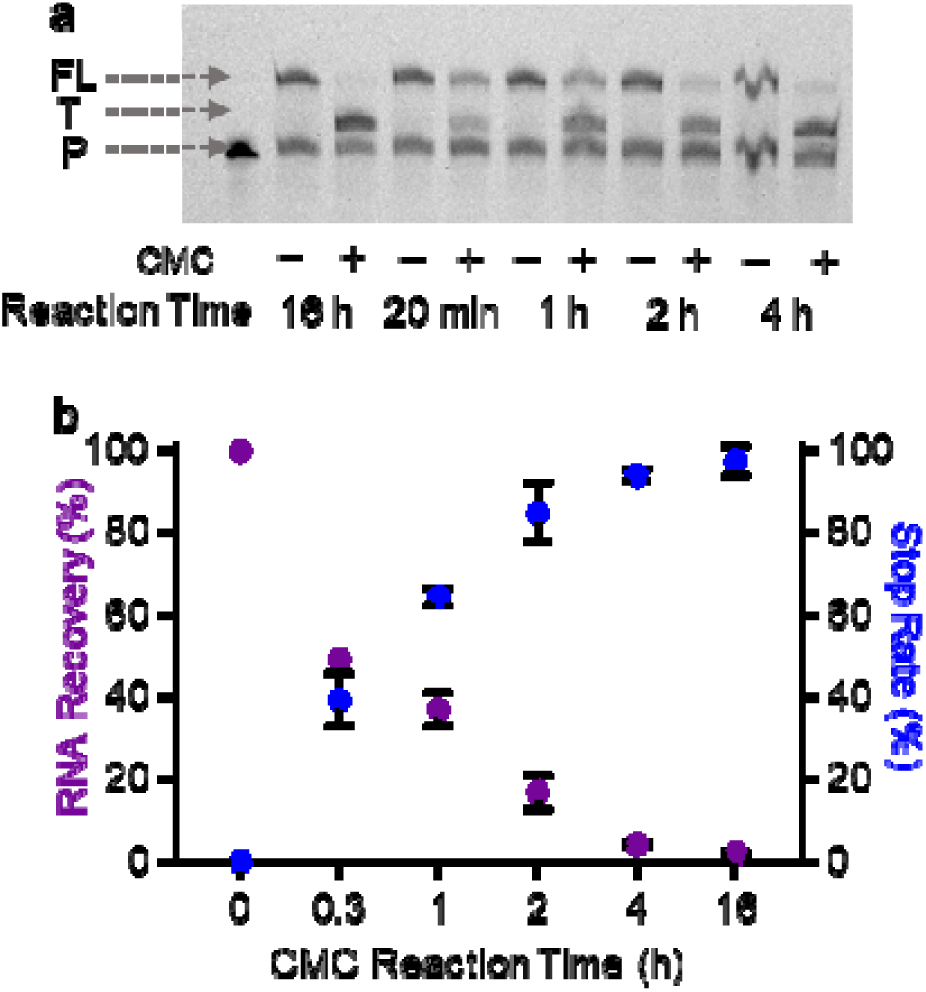
Characterization of Ψ into CMC-Ψ conversion and RNA loss by varying the duration of CMC reaction. **a**) Shown is the increasing Ψ into CMC-Ψ conversion efficiency of Ψ-oligo1 with the elongated CMC reaction duration, measured by the RT stop assay. **b**) Dependence of the Ψ into CMC-Ψ conversion of Ψ-oligo1 RNA and the RNA recovery after CMC treatment, upon the CMC reaction duration.

Here we measured the Ψ into CMC-Ψ conversion rates and RNA loss under various CMC reaction conditions by the SSIII RT stop assay. We used the RT stop rate as a proxy for estimating the Ψ into CMC-Ψ conversion (**Methods** and **Figure 3a**).^[23]^ With an increasing duration of the CMC reaction from 20 minutes to 16 hours, we observed increasing Ψ into CMC-Ψ conversion efficiency, accompanied with increasing loss of RNA (**Figure 3b**). CMC treatment for 4 or 16 hours achieved over 94% Ψ to CMC-Ψ conversion, however, these conditions resulted in over 95% loss of RNA. The loss of RNA can result from RNA degradation mediated by high concentration of CMC, and/or loss during the RNA purification procedure after the CMC reaction step. The 20-minute CMC condition (i.e., most frequently used in Ψ-sequencing methods as listed in **Figure S4b**) recovered 50% of the input RNA oligonucleotide; however, this condition showed only 40% conversion of Ψ to CMC-Ψ measured by the RT stop assay (**Figure 3** and **Figure S4c**). Interestingly, with 2-hour CMC treatment condition, the oligo RNA reaches ∼85% Ψ into CMC-Ψ conversion efficiency and was able to recover a decent amount (∼14%) of input RNA oligonucleotide after treatment (**Figure 3**). We reasoned that although elongated reaction time may reduce RNA recovery, it can potentially promote the detection sensitivity for Ψ due to more favorable Ψ to CMC-Ψ conversion. We decided to investigate how different CMC reaction conditions impact the performance of Ψ-seqs and proceeded testing the 20 minutes and 2 hours CMC treatment conditions by Ψ-seq.

### Profiling of RT signatures via “piloting” Ψ-seq libraries

To examine RT signatures by RT-1306 under various CMC treatment conditions by Ψ-seq, we prepared six “piloting” Ψ-seq libraries from total RNA extracted from HEK-293T cells, utilizing previously reported ligation-based library preparation protocols.^[8a, 8b, 21]^ These six “piloting” Ψ-seq libraries include libraries prepared by varying CMC reaction conditions and RTases (**Figure S5** and **Methods**), and sequenced with ∼50K reads per library at low cost (∼$50 per library). With these libraries, we primarily focused on assessing the RT signatures based on the reported Ψs within human rRNAs.^[22]^ The prepared libraries showed 54-58% alignment to the rRNA reference genome (**Figure S6**); encouragingly, RT-1306 generated mutation signatures at previously reported Ψs in the 18S and 28S rRNAs upon manual inspection. In contrast, wtRT and SSIII produced stop signatures with no detectable mutations (**Figure 4a**). Interestingly, the mutation rate at the Ψ1077 in 18S rRNA increased by ∼5-fold when the CMC reaction duration increased from 20 minutes to 2 hours (**Figure 4b**), which is consistent with the improved conversion of Ψ to CMC-Ψ observed on oligonucleotide RNAs (**Figure 3**).

**Figure 4.**
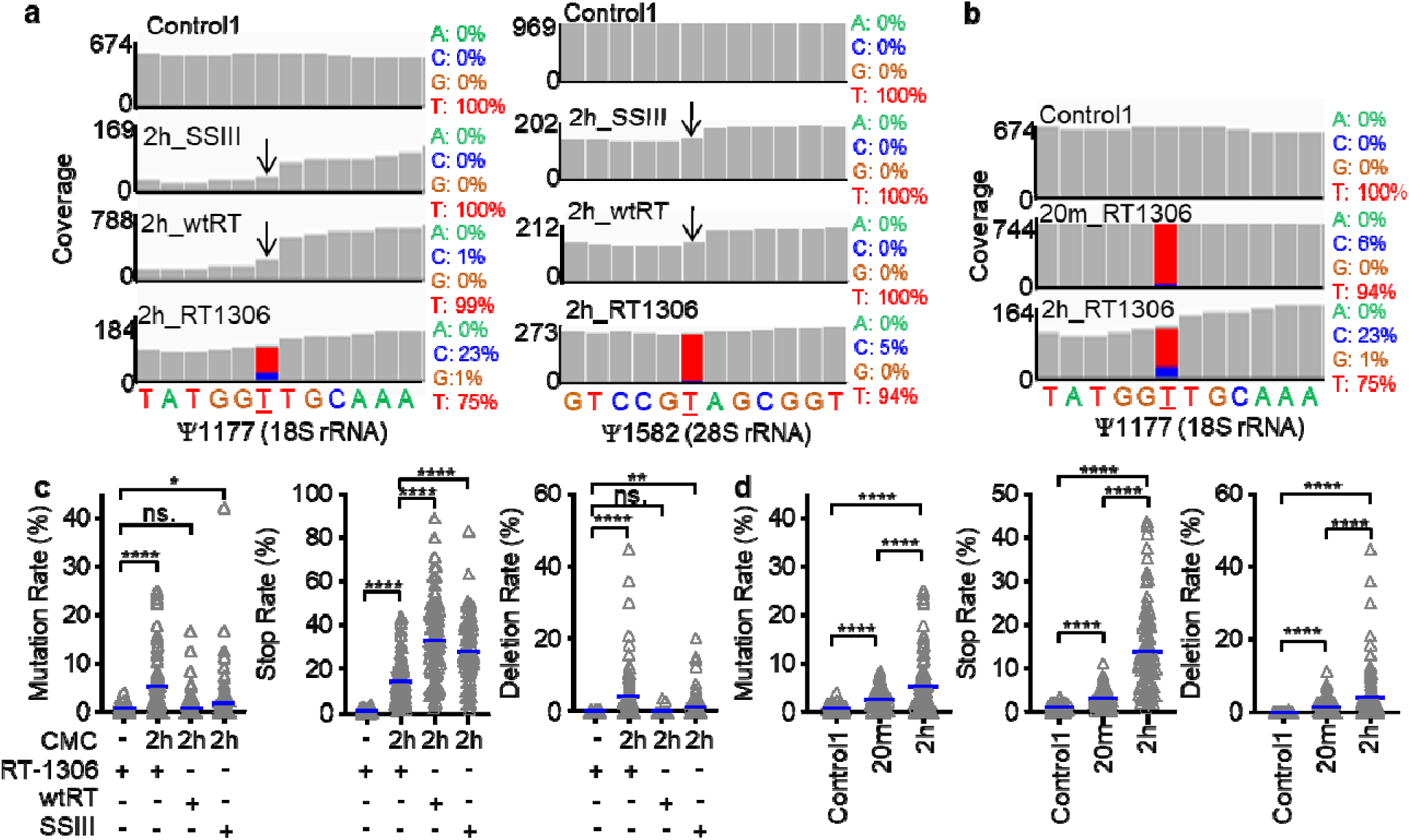
Promoted mutation signatures against CMC-Ψ in rRNAs by RT-1306 in “piloting” Ψ-seq libraries. Representative IGV views showing RT mutations at **a**) two example Ψ sites in 18S and 28S rRNAs. **b**) Ψ1177 in 18S rRNA with 20-minute or 2-hour CMC reaction. **c**) Profiling of RT mutation, stop and deletion signatures at all reported Ψ sites in rRNAs by RT-1306, with statistical comparisons to wtRT and SSIII. **d**) Profiling of RT signatures by RT-1306 at Ψ sites in rRNAs detected with 20-minute and 2-hour CMC reaction. *P* values were calculated by two-sided Student’s t-test, with the significance levels noted where ns. = not significant, \**P* < 0.05, \*\**P* < 0.01, \*\*\**P* < 0.001, \*\*\*\**P* < 0.0001.

Next, we examined all 105 documented Ψ sites in human 5.8S, 18S and 28S rRNAs (**Methods**).^[22]^ Strikingly, RT-1306 generated significantly promoted mutation signatures among Ψ sites with CMC treatment, showing ∼9-fold increase of the averaged mutation rate comparing with that of the “Control1” library (only OH^-^ treatment) (**Figure 4c** and **Figure S5a**). In contrast, wtRT and SSIII showed no or low extent of increase of mutation rates at Ψ sites (**Figure 4c**). Additionally, we analyzed all other possible RT signatures beyond mutation rates, including RT stops, deletions, and insertions (**Methods**). The RT signature profiling revealed that RT-1306 generated complex RT signatures against CMC-Ψ adduct in rRNAs, including mutation, stop and deletion, but not insertion. In contrast, both wtRT and SSIII generated predominantly RT stop signature (**Figure 4c** and **Figure S7a**). We observed significantly decreased RT stop rates against CMC-Ψs in rRNAs for RT-1306, comparing to those of wtRT and SSIII (**Figure 4c**), which is consistent with the improved read-through efficiency by RT-1306 against the CMC-Ψ RNA oligos (**Figure 2a**).

Next, we assessed the background noise level by profiling the RT signatures against 1088 unmodified U sites in 5S, 5.8S, 18S and 28S rRNAs.^[26]^ Not surprisingly, we found RT stops present much more prominent background noise relative to mutation, deletion and insertion signatures, only considering the non-CMC “Control1” library (**Figure S7b**). Upon 2-hour CMC treatment, all three RTases (SSIII, wtRT, and RT-1306) showed significantly increased RT stops; among the three RTases, the background RT stop is the least significant for RT-1306 (**Figure S7b**).

Interestingly, RT signatures against CMC-Ψ got promoted by 2-5-fold upon increasing the reaction duration of CMC treatment from 20 minutes to 2 hours, suggesting increased conversion ratio from Ψ to CMC-Ψ (**Figure 4d**). Importantly, the 2-hour CMC treatment did not increase the background mutation and deletion against unmodified Us by RT-1306 compared to the 20-minute reaction, though moderate increase was observed for the background stop signature (**Figure S7c**). In summary, the piloting Ψ-seq provided a low-cost and efficient method for validating library construction method, and profiling RT signatures on the abundant rRNAs with variable chemical treatment and RT conditions. We confirmed that mutation signature of RT-1306 against the CMC-Ψ was promising to map Ψs in biological RNAs and while the elongated reaction duration of CMC treatment can be promising to improve the detection sensitivity of Ψ, the level of background noise must be assessed accordingly.

### Development of Mut-Ψ-seq and quantitative assessment of Ψ-identification efficiency by ROC curve analyses

To apply RT-1306 into Ψ identification in coding and non-coding RNAs, we developed “Mut-Ψ-seq”. We treated fragmented polyA-enriched RNAs from HEK-293T cells (two biological replicates) with 20 minutes or 2 hours CMC reaction followed by alkaline treatment. The treated RNA was ligated with the 3’ adapter sequence and reverse transcribed by RT-1306. The cDNA product was ligated with the 5’ adapter followed by PCR amplification into NGS libraries (**Figure 5a**, **Table S1**, and **Methods**). To benchmark the ligation-based protocol, we prepared two libraries by 20 minutes CMC treatment condition and SSIII as previously reported (**Figure 5a**).^[8c]^ The resulted library samples (i.e. PCR products) were around 230 base pairs in size, except for the two libraries prepared from RNAs treated with 2-hour CMC reaction which showed significantly smaller library size (**Figure S8**). All libraries were subjected to deep sequencing with ∼40 million reads per library (**Methods**).

**Figure 5.**
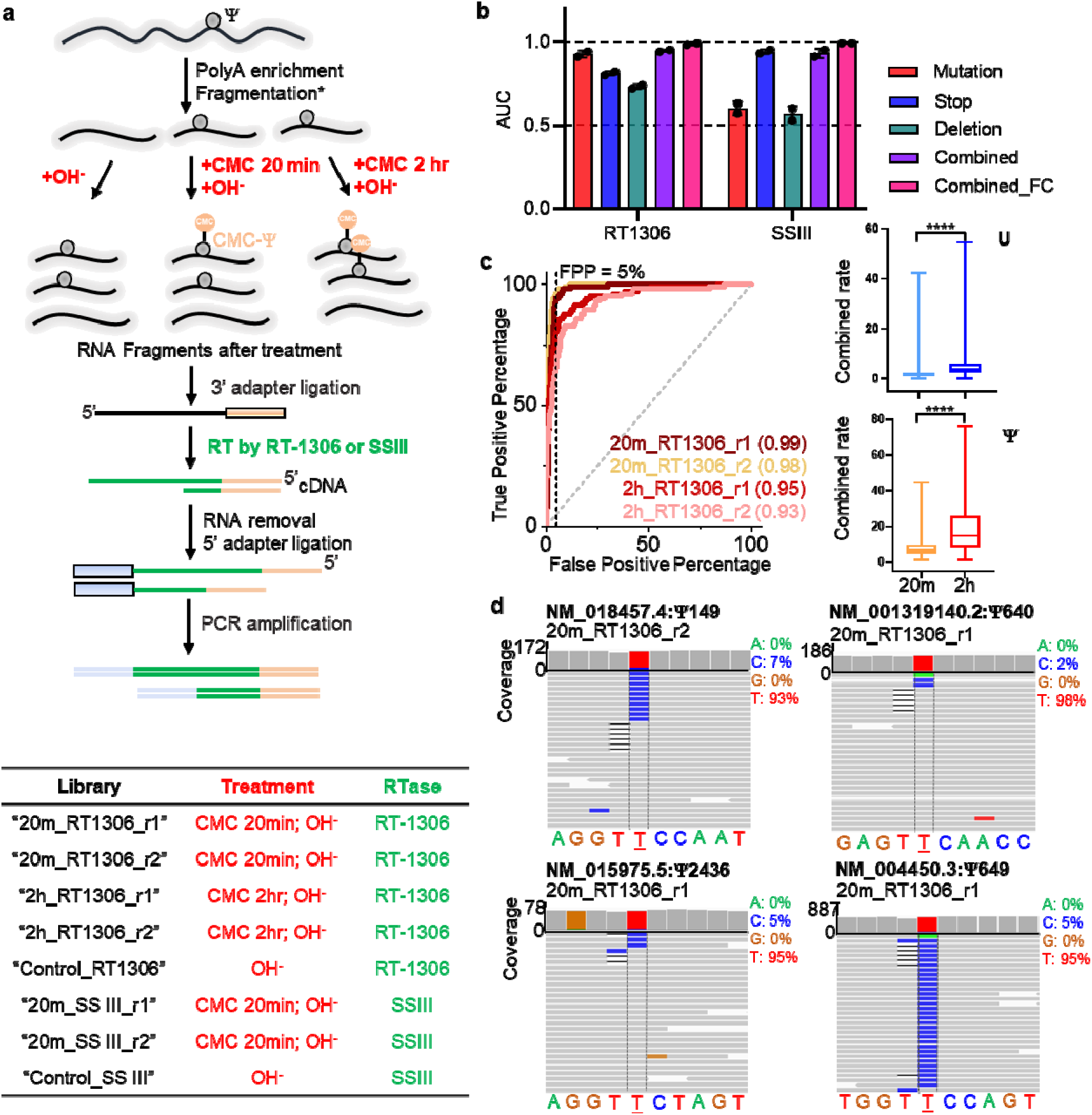
Development and results of Mut-Ψ-seq. **a**) Workflow and list of prepared libraries for Mut-Ψ-seq. **b**) Comparison of Ψ identification efficiency by individual or combined RT signatures generated by RT-1306 or SSIII, through the AUC values derived ROC curve analyses against reported Ψs and Us in rRNAs (**Figure S9** and **Figure S10**). **c**) Shown on the left is the ROC analyses for assessing Ψ identification efficiency of RT-1306 by 20-minute or 2-hour CMC reaction. Shown on the right are the distributions of observed combined rates generated by RT-1306 against reported 1088 Us (upper panel) and 105 Ψs (low panel) in rRNAs. *P* values were calculated by two-sided Student’s t-test, with the significance levels noted where \*\*\*\**P* < 0.0001. **d**) Representative IGV views of the RT mutation signature suggesting the occurrence Ψ in GUΨC sequence in four mRNAs.

We first assessed the Ψ-identification efficiency of the reported 105 Ψ and 1088 non-Ψ U sites in rRNAs^[26]^ by the ROC curve analysis and calculating the area under the curve (AUC). Briefly, AUC = 1 represents the perfect Ψ-identification sensitivity and specificity (*i.e.* no false positive or false negative discoveries), and AUC = 0.5 represents random selection of Ψs and unmodified Us.^[8a, 8b, 23]^ SSIII control libraries showed stop signatures at Ψs and the RT stop signatures robustly identified rRNA Ψs with AUC*^stop^_SSIII_* = 0.93 or 0.95 for the two biological replicates, which benchmarked the high quality of libraries prepared via the current protocol (**Figure 5b** and **Figure S9a**). Interestingly, under the same 20 minutes CMC treated condition, the RT mutation signature alone by RT-1306 can identify Ψ sites in rRNAs with AUC*^mut^_RT-1306_*= 0.94 or 0.91 for the two biological replicates, suggesting the mutation signatures by RT-1306 can be used to identify Ψ sites. In contrast, the same ROC analyses yielded AUC*^mut^_SSIII_* = 0.63 or 0.57 using the mutation signature generated by SSIII to identify Ψ sites, suggesting near random selection (**Figure 5b** and **Figure S9b**).

Given RT-1306 generated context-dependent RT signatures against CMC-Ψs (**Figure 4c**), we systematically assessed the performances of distinct RT signatures for Ψ identification via the ROC analysis. For RT-1306, mutation, stop, and deletion all showed identification power for Ψs with AUC > 0.7; among the three signatures, mutation is the most efficient signature given the highest AUC. In contrast, SSIII produced primarily the stop signature (**Figure 5b**). The combined rate (*i.e.* mutation rate + stop rate + deletion rate) by RT-1306 showed improved identification efficiency rather than single RT signature, where AUC*^com^* = 0.95 or 0.94 (**Figure 5b** and **Figure S9c**). To rule out background RT signatures (e.g. RT stops due to RNA secondary structures), we calculated the fold change of combined rates between the CMC treated and untreated libraries. The ROC analysis revealed that the fold change gave the most robust identification power for Ψs with AUC*^com_fc^_RT-1306_*= 0.99 or 0.98 (**Figure 5b** and **Figure S10**). Lastly, the 2-hour CMC treatment condition increased the magnitudes of the combined rates compared to the 20 minutes condition. However, the ROC analysis showed slightly decreased efficiency for Ψ identification, compared to the 20-minute treatment, due to elevated noise level on unmodified U sites (**Figure 5c, Figure S9** and **Figure S11**). Therefore, we continued calling Ψs in mRNAs and non-ribosomal non-coding RNAs (ncRNAs) using the fold change of combined rates between the CMC treated and untreated libraries.

### Ψ-identification in UU sequence contexts mRNAs via Mut-Ψ-seq

Mut-Ψ-seq libraries were aligned onto the hg38 RefSeq reference genome including mRNA and ncRNA genes (**Figure S12**). Us in mRNAs and ncRNAs are initially identified to be potential Ψ sites if they show (i) significant fold change of combined rates for CMC-treated library compared to the untreated library, (ii) significant combined rates above the background level, and (iii) at least 5 read counts for the sum of reads that contain mutation, stop, and deletion in both biological replicates (**Methods**, **Figure S12**, and **Supplementary Data**). The thresholds of the fold change of combined rates are determined by the ROC curve analysis at 5% false-positive discovery rate for the reported Ψs in rRNAs, where FC^Com^ = 2.4 or 7.8 are set as cut-offs for Ψ identification for the 20-minute or 2-hour CMC treated libraries, respectively (**Figure S10** and **Figure S12**). The thresholds of background of the combined rates were derived based on the observed combined rates for the U sites in rRNAs within the same CMC-treated library; we used 5% or 9.6% as the cut-offs for the combined rates for Ψ calling for the 20-minute or 2-hour CMC treated libraries, respectively (**Figures S11-12**). Initial Ψ sites were called under such criteria for both the 20-minute and 2-hour CMC treated libraries (**Figure S12** and **Table S2**). We then overlap these initially identified sites with those reported by PRAISE^[17]^ to circumvent limitations caused by side reactions by a single chemical treatment.

We report a list of 294 high-confidence Ψ-containing genes identified by orthogonal chemical treatments (*i.e.* CMC chemistry in this study and bisulfite treatment used by ‘PRAISE’^[17]^) (**Figure S13a**). With this list of genes, we performed gene ontology (GO) analysis, which revealed that Ψ-modified mRNA was significantly enriched in translation-related biological processes, especially enriched in multiple ribosomal proteins mRNAs (**Figure S13b** and **Table S3**). Since the expression of ribosomal proteins tend to be regulated in a concerted manner,^[27]^ it is worth pursuing whether pseudouridination can provide an additional layer of concerted regulation for ribosome biogenesis. While these genes were identified by both Mut-Ψ-seq and PRAISE, the identified Ψs do not always overlap with single-base precision.

When insisting on single-base-level overlap, 44 high-confidence Ψ sites are robustly identified in mRNAs and non-ribosomal ncRNAs by orthogonal chemistry treatments (**Table 1**). 34 out of the 44 sites show the identified Ψ in UU-containing sequence contexts. We observe mutation signatures in 21 out of the 34 sites, which help precisely identify the location of Ψ in the UU sequence contexts (**Table 1**). For examples, Ψ649 was unambiguously assigned at the second U in the GUUC motif by mutation signature in the 3’-UTR of the ERH mRNA (NM_004450.3) (**Figure 5d**), which was previously assigned to be a substrate site for TRUB1.^[14, 16-17]^ PRAISE reported the same location as “648-649” with ambiguity raised by the deletion signature. Similarly, we found three other mRNA sites corresponding to the TRUB1 substrate sequence GUΨC and mutation signatures robustly captured the presence of Ψ in the second U positions (**Figure 5d**, **Figure S14a** and **Table 1**). In addition, Mut-Ψ-seq identified a previously reported Ψ250 in the stem-loop 3 of the 7SK RNA written by the writer guided by the H/ACA box small RNA;^[28]^ the mutation signature called the nearby 247 site is also modified by Ψ in the AUΨUG sequence, which was also reported by PRAISE within an ambiguous sequence (“246-248”).^[17]^ Interestingly, the unmodified sequence of stem-loop 3 of 7SK was reported to show two-state conformational ensemble in solution; U250 occurs as an unpaired internal loop residue in one state, while pairs with A228 adjacent to an asymmetric loop in the other state.^[29]^ It would be interesting to assess whether and how the presence of Ψ at 250 and 247 modulate the structural dynamics of the stem-loop 3 and thus the regulatory function of the 7SK RNA.^[29-30]^

**Table 1.**
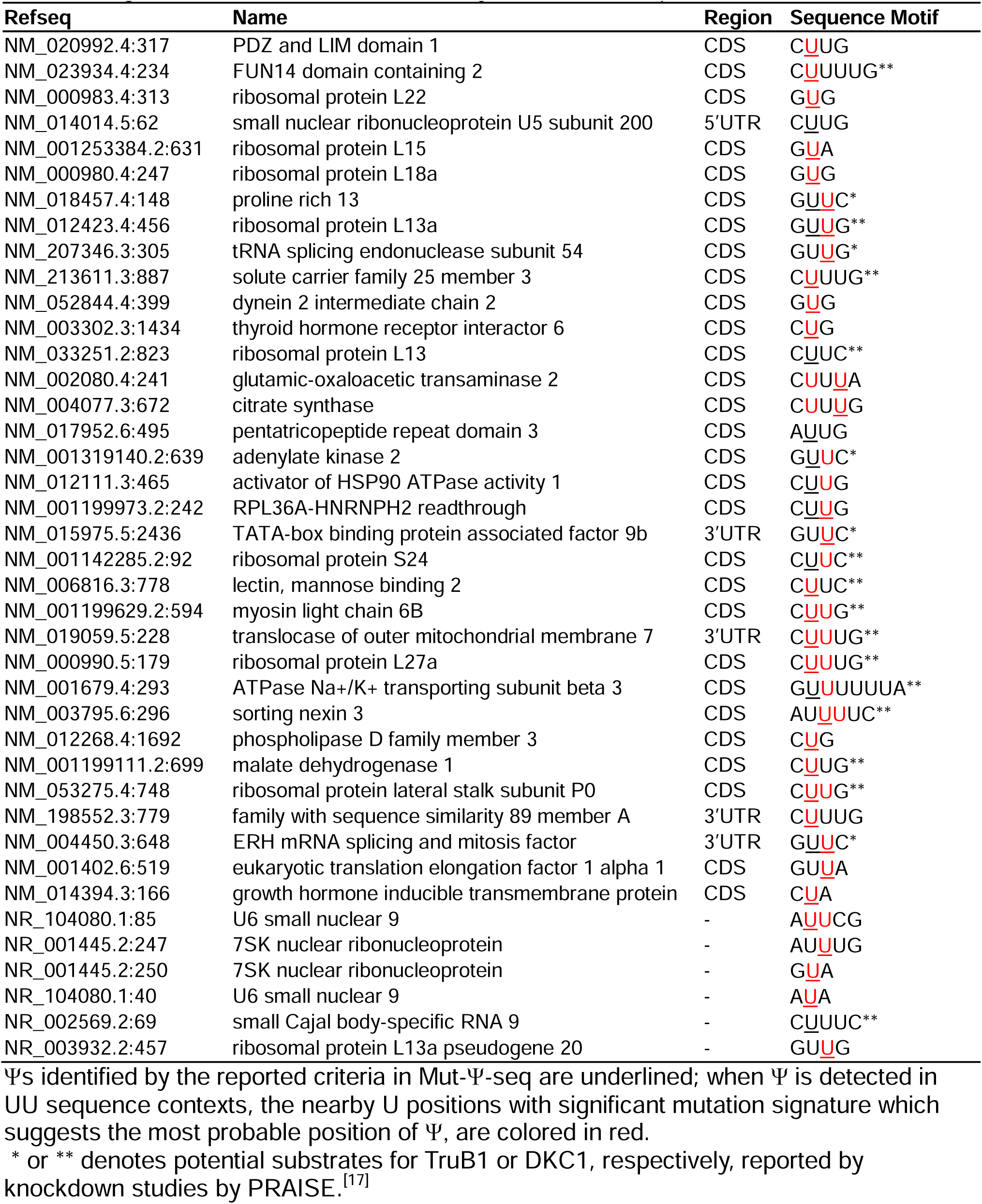
High-confidence Ψ sites identified by both Mut-Ψ-seq and PRAISE.

Notably, the initial assignment of Ψ sites in UU-containing sequences by the combined rates (mutation + stop + deletion) can be inaccurate by any prominent deletion rates (**Figure 5d**). The alignment tool in use (Bowtie 2^[31]^) aligns all deletions to the first U (from the 5’ end) in any sequences containing consecutive Us. This features a major limitation in using deletion signatures to identify Ψ in ‘UU’ sequences. Here we show the mutation signature against the classic CMC-treated biological RNAs can resolve the most probable Ψ sites in UU-containing sequences.

Mutation signatures revealed the diverse occurrence of Ψ in the UU-containing sequence contexts. In the high-confidence list of Ψ sites, Ψ was found at the second U in all eight GUUV (V = A/C/G) sequences. We found 6 mRNAs that contain one or two Ψ sites in the CUUG sequence context including two each of CΨUG, CUΨG, and CΨΨG. Only a subset of Us were found to be Ψ in short U-tract sequences such as CΨUUG, CΨΨUG, CΨUΨG, GUΨUU and AUΨΨU (**Table 1** and **Figure S14b**). Notably, writers for Ψ in UU-containing sequences have yet been established, except TRUB1 for GUΨC.^[14, 16-17, 32]^ “PRAISE” reported Ψs in a subset of UU-containing sequences can be regulated by DKC1 by Ψ mapping DKC1-knockdown (**Table 1**);^[17]^ however, the guide RNAs for the reported sites remain unclear. The precise locations of Ψ in these sequences are critical for the future identification of writers, especially beneficial for predicting guide RNA binding sites for RNA-dependent Ψ writing mechanisms in mRNAs and ncRNAs.

## CONCLUSION

Here we report a pseudouridine sequencing method “Mut-Ψ-seq” with promoted mutation signature generated by the evolved RTase “RT-1306” reading through CMC-Ψ adducts in CMC-treated biological RNA from HEK-293T cells. ROC curve analysis is a powerful method in quantitatively assessing the performance of chemical treatment and RT signatures for identifying modifications. We demonstrated that the RT-signatures by RT-1306 robustly identified previous documented Ψ sites in human rRNAs with AUC = 0.98. Our sequencing data showed an increased CMC reaction duration can magnify the detection sensitivity of Ψ, however it increased the level of background noise and RNA loss, and didn’t improve overall Ψ identification efficiency in Ψ-mapping efforts. Mut-Ψ-seq demonstrated great potential in identifying Ψ at base-resolution in UU-containing sequence contexts. The promoted mutation signatures generated by RT-1306 against the CMC-Ψ adduct enabled the determination of Ψ occurrence in UU-containing sequences in mRNA and non-ribosomal ncRNAs. We report high-confident lists of Ψ sites and genes identified by orthogonal mapping methods, which provide valuable insights for understanding the biogenesis and function of Ψ.

## METHODS

### DNA and RNA oligonucleotides

DNA primers, primers with FAM labeling and RNA Ψ-oligo1 for *in vitro* assays and cloning were ordered from Integrated DNA Technologies, Inc. (IDT), with standard desalting. RNA Ψ-oligo2 was ordered from Keck Biotechnology Resource with standard desalting. Ligation adaptors used in piloting Ψ-seq libraries and Mut-Ψ-seq were ordered from IDT with high-performance liquid chromatography (HPLC) purification. Sequences are reported in **Table S1**.

### Expression and purification of wtRT and RT-1306

The plasmid of wtRT and RT-1306 were transformed into *E. coli* BL21(DE3) respectively followed by culturing at 37 °C.^[21]^ The expression of RT protein was induced by adding 0.5 mM of isopropyl-β-D-thiogalactoside (IPTG) into 1 L of cell culture with 80 μM kanamycin when the OD600 reached 0.6-0.8. *E. coli* cells were cultured at 37 °C for 2 hours and then at 16 °C for 16 hours under shaking with 220 r.p.m. Cells were harvested and resuspended in 40 mL lysis buffer (50 mM Na_2_HPO_4_ and NaH_2_PO_4_, 0.5 M NaCl, pH 7.8, dissolved half-tablet of the proteinase inhibitor cocktail, Pierce). The cells were then lysed by sonication and centrifuged at 12,000 r.p.m. for 40 minutes at 4 °C. Solubilized proteins in the supernatant were first purified using His60 Ni Superflow Resins (Takara Bio USA, 635660) and were eluted with 50 mM to 0.5 M gradient imidazole buffer containing 50 mM Na_2_HPO_4_ and NaH_2_PO_4_, pH = 6.0, 0.3 M NaCl and 10% glycerol. The eluted protein fractions were run through a desalting column (PD-10, GE Healthcare), the buffer was exchanged into 3 mL ion-exchange Buffer A (50 mM Bis-tris pH 7.0) with an additional 50 mM NaCl. Then, the fractions were subjected to Mono Q ion-exchange chromatography, where the protein was injected onto the column flushing with 97.5% Buffer A and 2.5% Buffer B (50 mM Bis-tris pH 7.0, 1 M NaCl) and the protein was recovered in the flow-through portion. The ion-exchange purification was found to be essential for effectively removing nuclease contamination. All purification steps were carried out at 4 °C or on ice. Fractions containing the expressed protein were combined and concentrated to 2.5 mL with a 30-kDa cut-off centrifugal filter (Millipore), run through the desalting column again, and eluted with the storage buffer (50 mM Tris-HCl, 25 mM NaCl, 1 mM EDTA, 50% glycerol, pH 7.0). Purified proteins were concentrated to 200-300 μL using a 30-kDa cut-off centrifugal filter, aliquoted, flash frozen in liquid nitrogen and stored at −80 °C.

### CMC labeling of RNA oligonucleotides and RNA recovery quantification

Synthetic RNA molecules were used in this study: Ψ-oligo1 and Ψ-oligo2 (**Table S1**). 10 μM RNA oligos were reacted with 0.2 M CMC (Chem-Impex) in BEU buffer (7 M urea, 4 mM EDTA, 50 mM Bicine, pH 8.5) at 37 °C for specific time (20 min, 1 hour, 2 hours, 4 hours, or 16 hours) followed by Oligo Clean & Concentrator Kits (OCC, Zymo Research, D4061) cleanup, eluted with 20 μL RNase free water. Purified RNA was then mixed with 2 volumes of sodium carbonate buffer (50 mM Na_2_CO_3_, 2 mM EDTA, pH 10.4) and incubated at 37 °C for 4 hours to remove CMC from Us and Gs, and purified with the OCC for subsequent characterizations. The concentration of eluted RNA was quantified through A260 reading measured by Nanodrop. RNA recovery was calculated by dividing the recovered RNA amount after 2-step treatment by the starting RNA amount, molecular weight change during this process is neglected.

### RT stop assay

The RT stop assays were performed in 10-μL reaction volumes containing 1× RT buffer, 4 pmol RNA substrates (with or without CMC treatment) and 4 pmol DNA primer with 5’-FAM label (**Table S1**), 1 mM of each dNTP and 2 μM purified RTase. The RNA substrate and DNA primer were added first and incubate in a thermocycler at 65 °C for 4 minutes, then 55 °C for 2 minutes, 45 °C for 2 minutes, and 37 °C for 2 minutes for annealing. The RT reactions were then carried out at 37 °C for 1 hour followed by heating at 80 °C for 10 minutes to inactivate the RT (for SSIII, Invitrogen, 18080093, incubate at 25 °C for 4 minutes then 42 °C for 10 minutes, 52 °C for 40 minutes followed by heating at 70 °C for 10 minutes to inactivate the RT). Remove the RNA by adding 1 μL 1M NaOH, and incubating in a thermocycler at 95 °C for 15 minutes. 5 μL of the reaction was mixed with 5 μL 2× RNA loading dye and heated to 95 °C for 5 minutes. 7 μL of that mixture was then immediately loaded onto 15% denaturing 8 M Urea PAGE gel. Gel was pre-run for 30 minutes at 200 V, and continued running for 1 hour after sample loading. The gel was then imaged by fluorescence detection on a Bio-Rad ChemiDoc System and analyzed with Image Lab software. The gel was stained with 1x SYBR Gold (Invitrogen, S11494) in TBE buffer for 30 minutes, then the stained gel was then imaged by SYBR Gold detection on a Bio-Rad ChemiDoc System and analyzed with Image Lab software.

### Colony sequencing assay

The CMC reactions were performed as described above 10 pmol of CMC treated or untreated Ψ-oligo1 and Ψ-oligo2 RNAs were subjected to 20 μL of RT reactions containing 0.5 mM each dNTP, 20 U SUPERase·In RNase Inhibitor (Invitrogen, AM2696), 1 uM RT-1306, 75 mM KCl, 2 mM MgCl_2_, 50 mM Tris-HCl (pH 8.3); 10 μL of the RT reaction was used as the template for PCR reactions by adding 1 µL 10 mM dNTP Solution Mix, 0.5 µL Q5 High-Fidelity DNA Polymerase (NEB, M0491S), 10 µL 5 X Q5 reaction buffer, 23.5 µL water with 2.5 µL 10 µM forward and reverse primers (**Table S1**). PCR reaction was performed according to the manufacturer’s manual, and the products were purified by agarose gel purification. The PCR product was cloned into plasmid by Gibson Assembly and individual clone was picked and sent out for Sanger sequencing (**Figure S3a**). 10 and 7 colonies were picked for CMC treated Ψ-oligo1 and Ψ-oligo2 RNAs, respectively (**Figure 2b** and **Figure S3b**).

### Cell culture and total RNA extraction

HEK293T cells were maintained on 100 mm Surface Treated Tissue Culture Dishes (Fisherbrand, FB012924) in DMEM medium (Gibco, 10569010) supplemented with 10% FBS (Gibco) and 1% penicillin/streptomycin (Gibco, 15140148). The cells were maintained at 37 °C under a humidified atmosphere containing 5% CO_2_. Total RNA was extracted with TRIzol (Invitrogen, 15596018) followed by isopropanol precipitation, according to the manufacturer’s instructions, each plate of cells resulted ∼400 μg of total RNA. The resulting total RNA was treated with DNase I (NEB, M0303L) to avoid DNA contamination.

### RNA preparation for piloting Ψ-seq libraries

27.4 μg of total RNA was fragmented into ∼150-200 nt fragments at 94 °C for 5 minutes using the magnesium RNA fragmentation buffer (NEB, E6186AVIAL), followed by purification with OCC, and eluted with 30 μL RNase free water (710 ng/μL). Into 3 μL fragmented RNA, added 10 μL water and 10 μL 10 mM EDTA, heated at 80 °C for 5 minutes, and immediately placed on ice. 20 μL denatured RNA was added to 20 μL 0.4 M CMC in BEU buffer as the "CMC+" sample; or added to 20 μL BEU buffer without CMC as the "Control" sample. The CMC reaction was carried out at 37 °C for specific time (20 minutes, 2 hours, **Figure S5**).

This step was then followed by purification with OCC eluted in 15 μL water. RNA recovered from the CMC_20 min, CMC_2h and control samples were next separately dissolved in 30 μL Na_2_CO_3_ buffer (50 mM sodium carbonate/sodium bicarbonate, pH 10.4, 2 mM EDTA) and incubated at 37 °C for 4 hours. An additional purification with OCC was then performed to recover the RNA, eluted with 15 μL RNase free water, and the RNA concentration is around 130 ng/μL.

### RNA preparation for Mut-Ψ-seq

Two biological replicates were prepared by harvesting cells from two plates at the same passage. For polyA+ RNA isolation, 75 μg of total RNA were subjected to two sequential rounds of polyA^+^ selection for each biological replicate using oligo(dT)_25_ Dynabeads (Invitrogen, 61005) according to the manufacturer’s instructions. 700 ng polyA+ RNA was fragmented into ∼150-200 nt fragments at 94 °C for 3 min using the magnesium RNA fragmentation buffer, followed by purification with OCC. We shorten incubation time due to the possibility that CMC would digest RNA to some degree. Then the fragmented RNA was treated with CMC for 20 minutes or 2 hours (**Figure 5**), followed by base treatment. Detailed procedure is as same as described for RNA preparation for piloting Ψ-seq libraries. Finally, the RNA eluted with 18 μL RNase free water.

### Library construction

The library was prepared following our previously reported procedure with slight changes.^[21]^ 3′-End repair and 5′-phosphorylation were conducted with T4 polynucleotide kinase (PNK) (NEB, M0201S). 16 µL RNA was mixed with 2 µL 10× T4 PNK reaction buffer and 1 µL T4 PNK, 1 μL SUPERase•In RNase Inhibitor, and incubated at 37 °C for 1 h; followed by RNA Clean and Concentrator (Zymo Research, R1017) purification eluting with 10 µL RNase free water. To perform 3′-adapter ligation, 10 µL 3′-repaired and 5′-phosphorylated RNA fragments were incubated with 2 µL 10 µM RNA 3′ Adapter (5′App-NNNNNATCACGAGATCGGAAGAGCACACGTCT-3SpC3) at 70 °C for 2 minutes and placed immediately on ice. Then, 2 µL 10× T4 RNA Ligase Reaction Buffer (NEB), 6 µL PEG8000 (50%), 1 µL SUPERase•In RNase Inhibitor and 1 µL T4 RNA Ligase 2 truncated KQ (NEB, M0373L) were added to the RNA-adapter mixture. The reaction was incubated at 25 °C for 2 hours followed by 16 °C for 12 hours. 1 µL 5′-deadenylase (NEB, M0331S) was added into each ligation mixture by incubation at 30 °C for 1hour followed by adding 1 µL RecJf (NEB, M0264L) for ssDNA digestion at 37°C for 1 hour. Add 1 µL Proteinase K (Invitrogen, EO0491) 37°C for 15 minutes. Bring reactions to 50 µL by adding 27 µL RNase free water and perform OCC purification, the 3′-end-ligated RNA was extracted by OCC and eluted with 12 µL RNase free water. 1 µL RT primer was added to RNA and incubated in a thermocycler at 65 °C for 4 minutes, then 55 °C for 2 minutes, 45 °C for 2 minutes, and 37 °C for 2 minutes for annealing. For reverse transcription with HIV reverse transcriptase, to this was added 5X RT buffer, 1 µL 10 mM dNTP Solution Mix, 1 µL SUPERase•In RNase Inhibitor and 2 µL 10 µM RT-1306 or wtRT. The reaction was mixed well and incubated at 37 °C for 1 hour, was then heated at 80°C for 5 minutes. For reverse transcription with SSIII, 4 µL 5× first strand buffer, 1 µL 10 mM dNTP Solution Mix, 1 µL 100 mM dithiothreitol, perform RT at 25 °C for 4 minutes then 42 °C for 10 minutes, 52°C for 40 minutes followed by heating at 70 °C for 10 minutes. cDNA clean up following OCC instructions with 7 µL water. Add 0.8 μL 80 μM 5’ adaptor (5’ Phos-NNNNNAGATCGGAAGAGCACACGTCTG-3SpC3) and 1 μL DMSO into 5.5 μL cDNA sample, and mix well. Heat at 70°C for 2 minutes and then chill on ice immediately. Then 2 μL 10x RNA ligation buffer (NEB), 0.2 μL 100 mM ATP, 9 μL 50% PEG8000, 1.5 μL Rnl1 (high conc) ligase (NEB, M0437M) were added the ligation mixture was incubated at 25 °C for 12 hours. Add 1 μL Protease K to all reactions and incubate at 37 °C for 15 minutes. Add 29 μL RNase free water to bring reaction volume up to 50 μL and add 350 μL DNA binding buffer in the kit, perform DNA Clean and Concentrator (Zymo Research, D4004) using 1:7 reaction to binding buffer ratio to clean up and elute with 20 μL total volume (10 μL each, two times). 8 µL eluted cDNA was used for each 13-cycle PCR amplification reaction, which was performed with the NEBNext Universal PCR Primer for Illumina (NEB) and indexed primers (NEB). All libraries were purified on a 3% low melting point agarose gel.

### Sequencing data processing

The sequencing data (the R1 reads of the pair-end data) were subjected to deduplication by the BBMap tool “clumpify” (v.38.73)^[33]^ with the option ‘dedupe subs = 0’ to remove PCR duplicates. Adaptors were then trimmed by the cutadapt tool (v.1.15)^[34]^ while reads were filtered by quality and length with options ‘-a AGATCGGAAGAGCACACGTCTGAACTCCAGTC -q 20 -m 30’. Processed reads were aligned to the human rRNA genes or RefSeq reference transcriptome (GRCh38) using and Bowtie 2 (v.2.4.0)^[31]^ with parameter --very-sensitive-local (**Figure S9a** and **Figure S12**). Read counts for individual base types, deletions, and insertions at each base position were counted by the ‘bam-readcount’ tool^[35]^ in reference to the script ‘bam-readcount.sh’; the output of bam-readcount were further parsed by an in-house python script “parse_R1_with_indel-r.py” or “parse_R1_mut_del.py” to calculate the following rates(**Supplementary Data**). At each U position, mutation, deletion, and insertion rates are calculated as following:

Mutation rate = (A-readcount + C-readcount + G-readcount) / total-readcount;

Deletion rate = deletion-readcount / total-readcount;

Insertion rate = insertion-readcount / total-readcount.

When the RTase stops at the Ψ-CMC adduct, the cDNA terminated significantly at the nucleotide 3’-adjacent to the Ψ.^[8a, 8b]^ To quantify RT stop rates at each base position (e.g. the ***i*** nucleotide position), we used “bedtools genomecov -d -3” to count the number reads of which the 3’ ends aligned at the ***i***+1 position (*i.e.* readcount^3p^), and “bedtools genomecov -d” to count the total number of reads that aligned to the ***i***+1 position (readcount^total^). The stop rate at the position ***i*** is calculated by “Stop rate [***i***] = readcount^3p^[***i***+1]/readcount^total^[***i***+1]” (**Figure S9a**, **Figure S12a** and **Supplementary Data**).^[8b]^

### Calling Ψ sites

Combined rates were calculated by the sum of stop rate, deletion rate and mutation rate. Combined rate fold change was calculated using the equation Fold-change^Com^ = Combined rates^CMC+^/combined rate^Control.^ We detect a position as Ψ only when the following criteria were met: (i) the sum of reads aligned to the U position containing mutation, stop or deletion must be no less than 5 in the CMC treated libraries; (ii) the combined rate for the position is greater than the maximum value of the combined rates (mean+ standard deviation) of U determined by histogram analysis of combine rates in rRNA; (iii) the combined rate fold change is determined by the ROC. The combined rate fold change should be greater than the threshold value of which the false positive rate is less than 5%.

### Gene ontology analysis

The Gene Ontology (GO) analysis was performed using the Metascape^[36]^ bioinformatics database with default settings (https://metascape.org/).

### Statistical analysis

Statistical analysis was performed using GraphPad Prism 9.5.0 (GraphPad Software, Inc.). Asterisks denote statistical significance (ns, not significant; *, *P* < 0.05; **, *P* < 0.01; ***, *P* < 0.001; ****, *P* < 0.0001).

## Data availability

Raw and processed piloting Ψ-seq and Mut-Ψ-seq data are available at NCBI Gene Expression Omnibus, accession number GSE269406. The plasmid for bacterial expression of RT-1306 is available on Addgene with the ID 131521. The data that support the findings of this study are available from the corresponding author upon request.

## Code availability

Processing scripts for piloting Ψ-seq library and Mut-Ψ-seq and the scripts descriptions are available in the **Supplementary Data**.

## ACKNOWLEDGEMENT

We thank the Office of the Vice Provost for Research at Boston College for providing the Bio-Rad ChemiDoc System used for the research, and the BC Research Services for providing the Linux cluster in support of the bioinformatic analysis. We thank the laboratory of Prof. Welkin Johnson at Boston College for sharing the HEK293T cell line.

## SUPPLEMENTARY INFORMATION

Supplementary figures and description of processing scripts.

## SUPPLEMENTARY DATA

Supplementary tables and data processing scripts.

## Entry for the Table of Contents

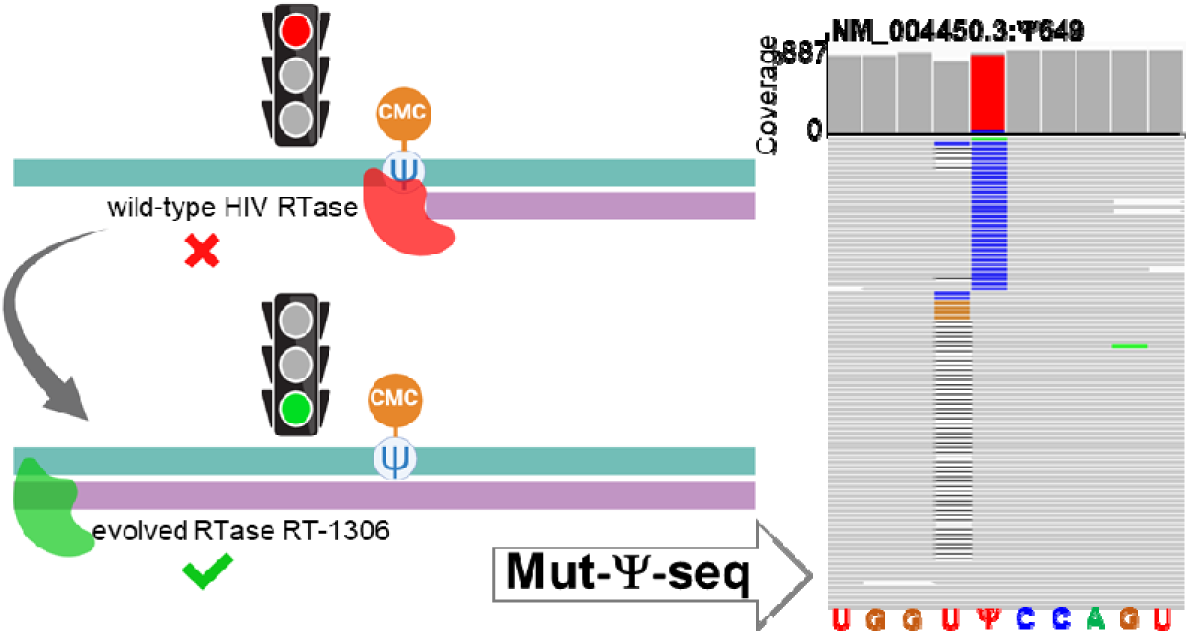

Conventional pseudouridine (Ψ) detection methods rely on reverse transcription truncation at CMC-Ψ. However, this study developed an evolved reverse transcriptase capable of reading through the bulky adduct and introducing a misincorporation at Ψ sites. This mutation signature enables the determination of Ψ positions within consecutive uridine sequences.

