## Supplementary material for "Promoted Read-through and Mutation Against Pseudouridine-CMC by an Evolved Reverse Transcriptase": SI

**Table of Contents**

1. Supplementary Figures
2. Description of Processing Scripts
3. Description of the Supplementary Tables

1. **Supplementary Figures**

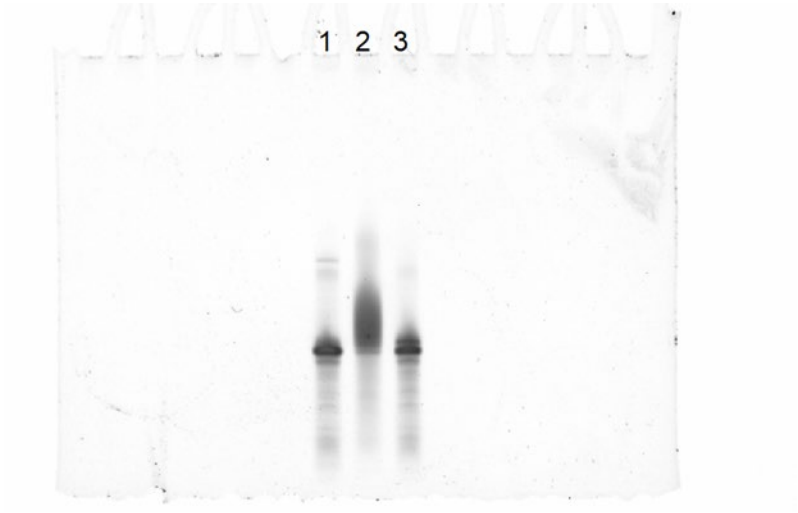


**Figure S1**. Electrophoretic analysis of CMC reaction with model RNA oligo, Ψ-oligo1 by 15% 8M Urea PAGE gel. Lane 1: Ψ-oligo1, lane 2: Ψ-oligo1+CMC, lane 2: Ψ-oligo1+CMC+OH-.

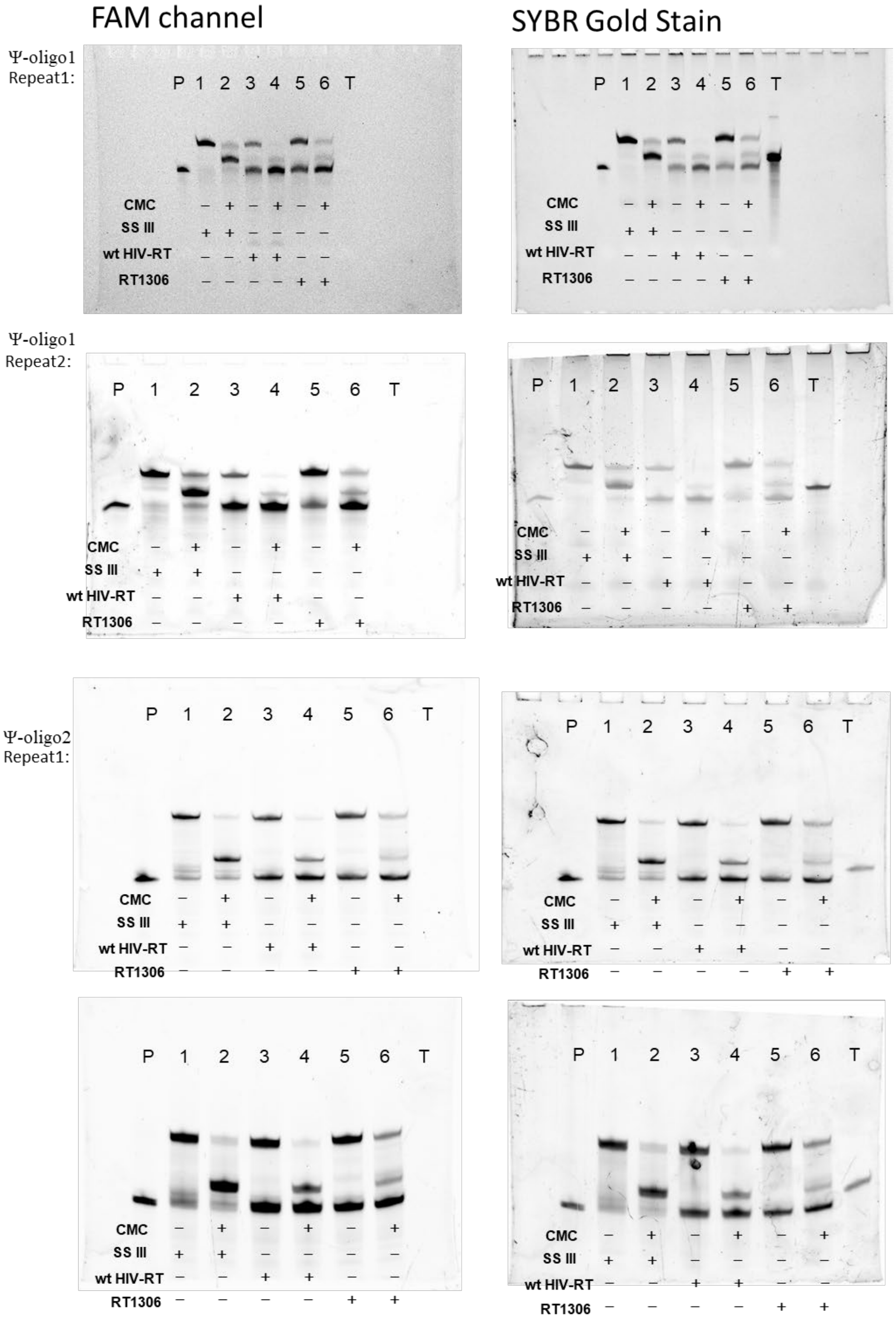


**Figure S2.** Electrophoresis gel showing three RTases RT-stop assay results with a 33-mer Ψ-oligo1 and 40-mer Ψ-oligo2 with (“+”) or without (“-“) CMC treatment.

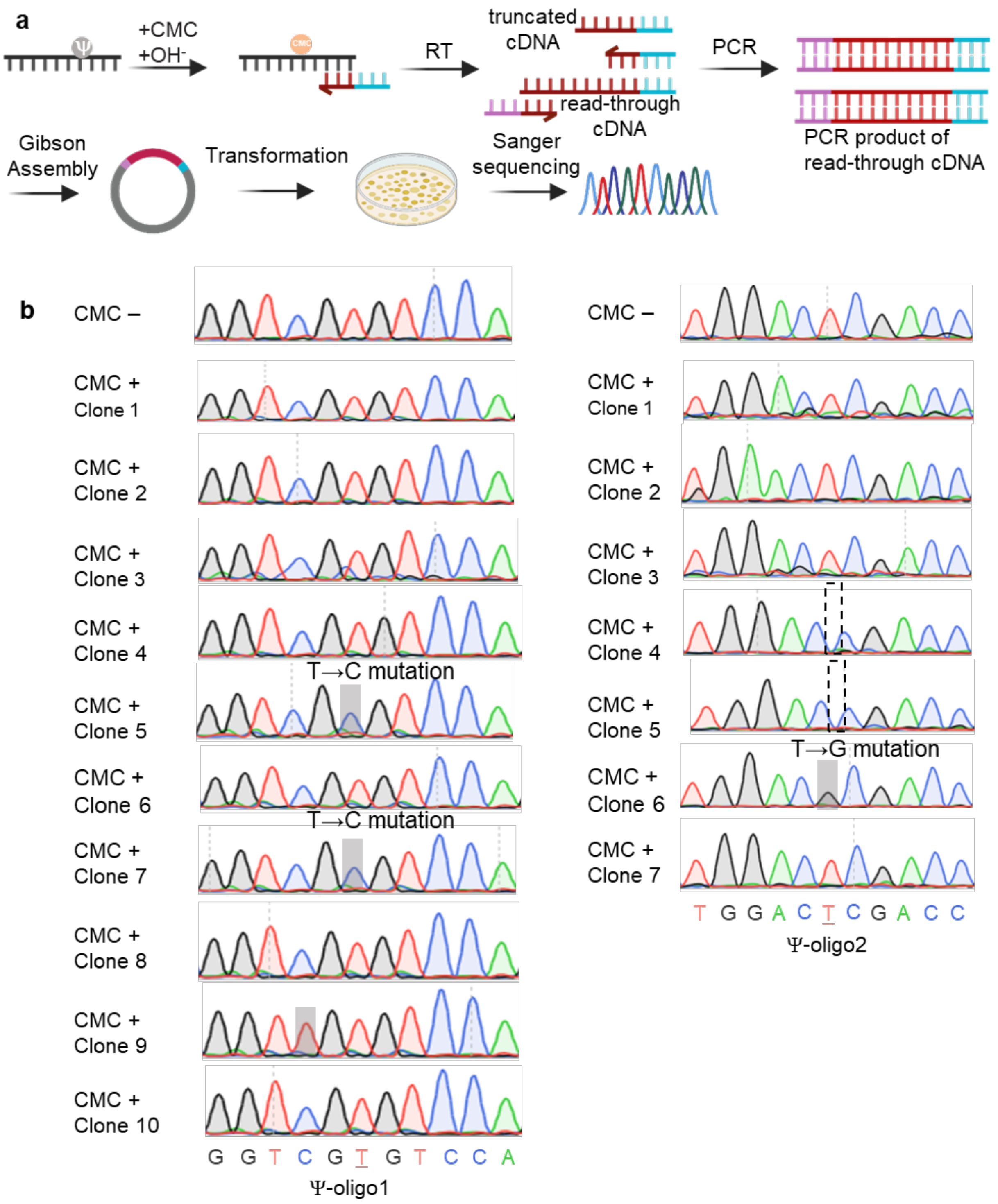


**Figure S3.** a) Scheme of colony sequencing. CMC-treated RNAs were reverse transcribed into cDNA, amplified by PCR and cloned into plasmids. Individual colonies were picked for Sanger sequence to read out cDNA sequences. b) Sanger sequencing of colonies prepared from the cDNA products by the RT-1306 of Ψ RNA oligos with and without CMC treatment.

**a**

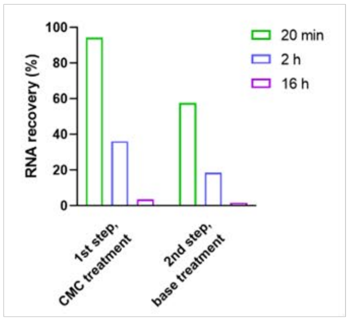


**b**

| reference | CMC treatment | Base treatment |
|---|---|---|
| Carlile *et al* Nature 2014 | 0.2 M or 0.4 M CMC, 40°C for 30 min | 50°C for 2 hr |
| Schwartz *et al* Cell 2014 | 0.17 M CMC, 37°C for 20 min | 37°C for 3 hr |
| Lovejoy *et al* PLOS ONE 2014 | 0.2 M CMC, 37°C for 20 min | 37°C for 4 hr |
| Li *et al* Nature Chemical Biology 2015 | 0.2 M CMC-$N_3$, 37°C for 20 min | 37°C for 6 hr |

**c**

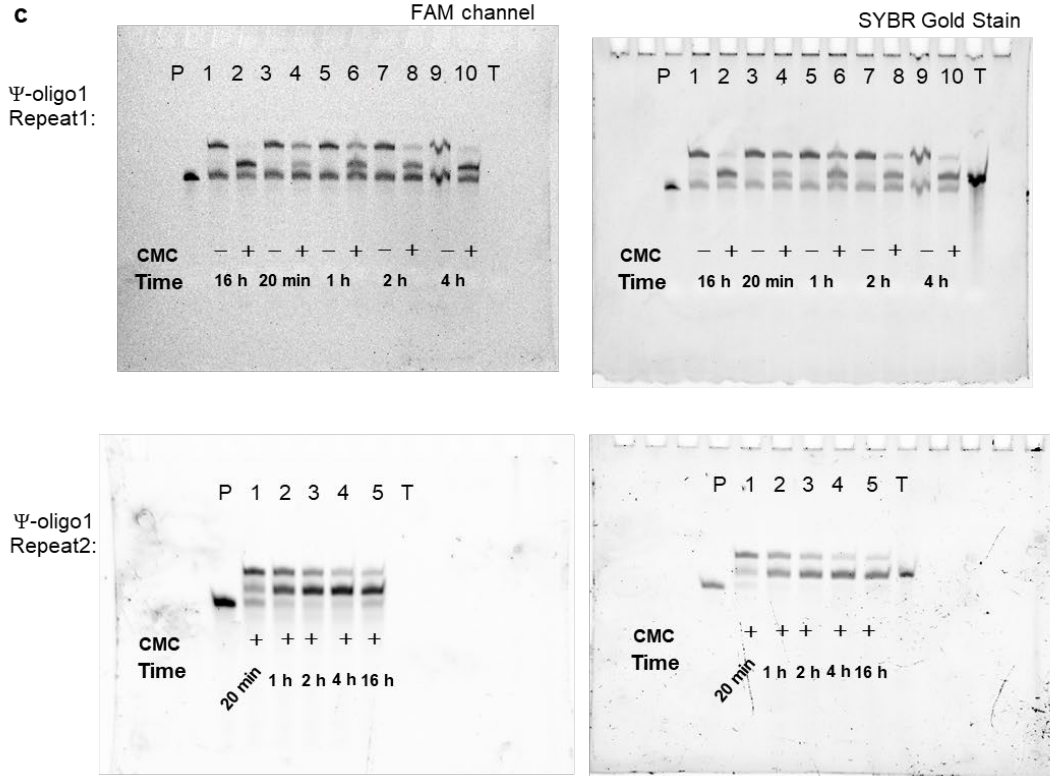


**Figure S4**. a) RNA loss by CMC and base treatment. b) Reported reaction conditions for pseudouridine sequencing. c) Characterization of Ψ to CMC-Ψ conversion and RNA loss under different CMC reaction durations.

a

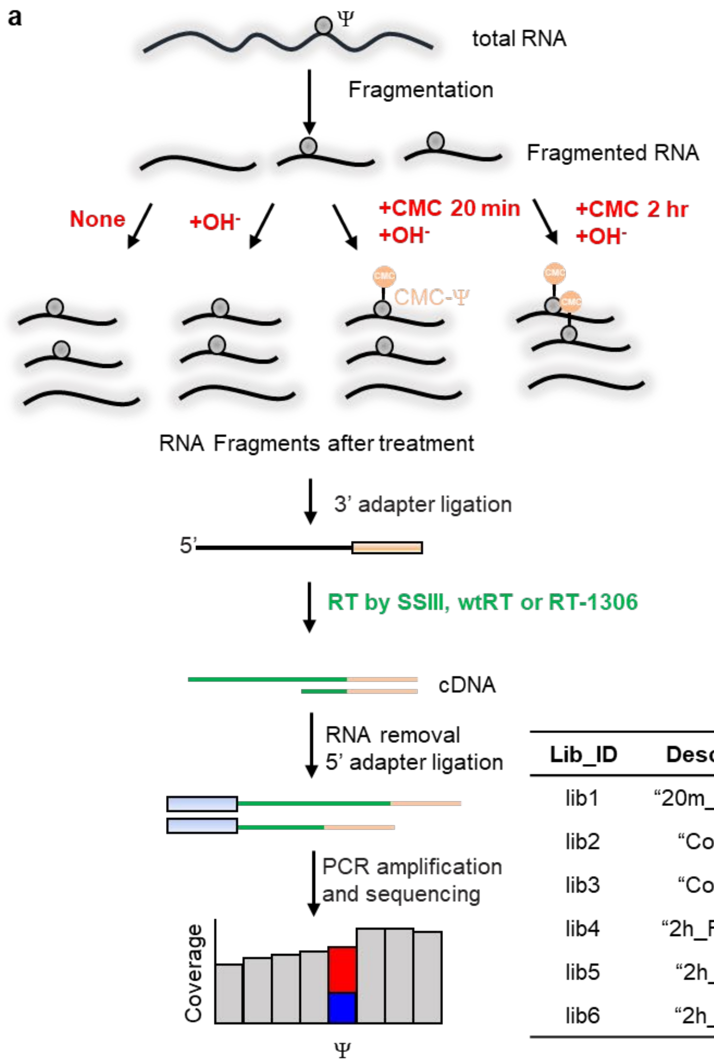


| Lib_ID | Description | Treatment | RTase |
| --- | --- | --- | --- |
| lib1 | "20m_RT1306" | CMC 20 min; $OH^-$ | RT-1306 |
| lib2 | "Control1" | $OH^-$ | RT-1306 |
| lib3 | "Control2" | None | RT-1306 |
| lib4 | "2h_RT1306" | CMC 2 hr; $OH^-$ | RT-1306 |
| lib5 | "2h_wtRT" | CMC 2 hr; $OH^-$ | wtRT |
| lib6 | "2h_SS III" | CMC 2 hr; $OH^-$ | SS III |

b

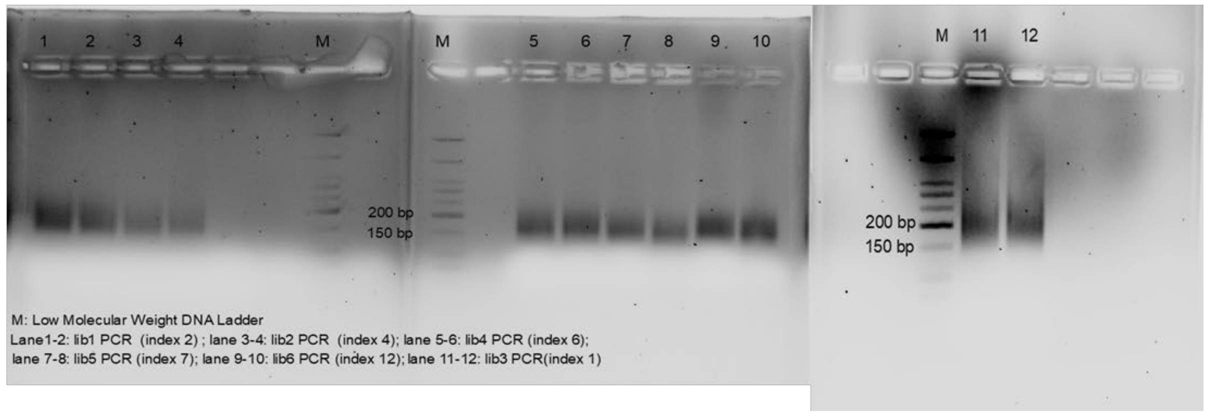


**Figure S5.** a) Flowchart for library preparation steps for the piloting Ψ-seq libraries. b) Gel data for characterize the sizes of the Ψ-seq libraries from CMC treated RNA or untreated control RNAs.

| Lib_ID | Description | Treatment | RTase | Alignment Rate to rRNA |
|---|---|---|---|---|
| lib1 | "20m_RT1306" | CMC 20 min; $OH^-$ | RT-1306 | 57.7% |
| lib2 | "Control1" | $OH^-$ | RT-1306 | 58.5% |
| lib3 | "Control2" | None | RT-1306 | 54.3% |
| lib4 | "2h_RT1306" | CMC 2 hr; $OH^-$ | RT-1306 | 55.2% |
| lib5 | "2h_wtRT" | CMC 2 hr; $OH^-$ | wtRT | 57.7% |
| lib6 | "2h_SS III" | CMC 2 hr; $OH^-$ | SS III | 57.4% |

**Figure S6.** Alignment rate for the piloting Ψ-seq libraries.

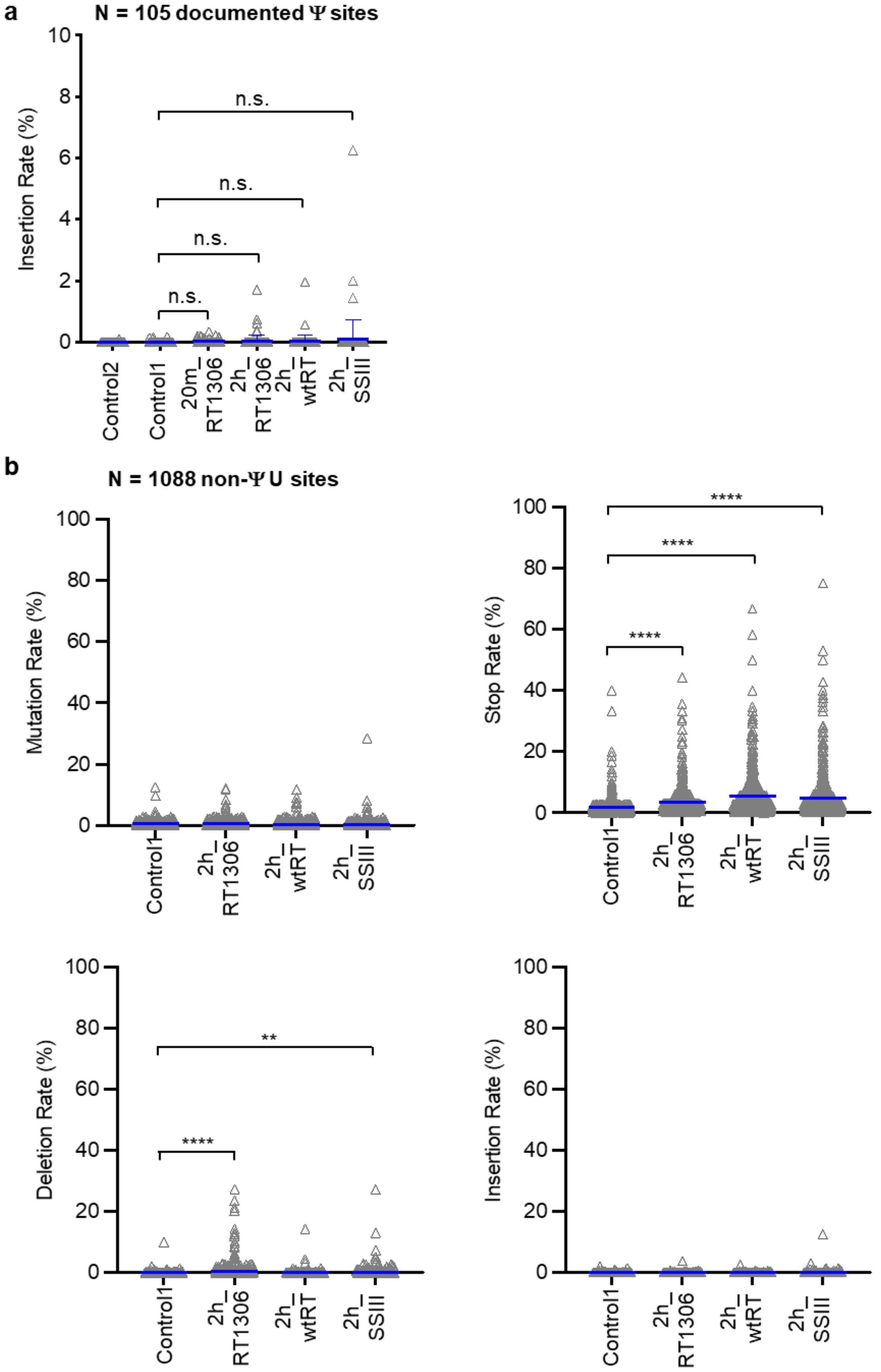
a
N = 105 documented Ψ sites
Insertion Rate (%)
10
8
6
4
2
0
n.s.
n.s.
n.s.
n.s.
Control2
Control1
20m_ RT1306
2h_ RT1306
2h_ wtRT
2h_ SSIII
b
N = 1088 non-Ψ U sites
Mutation Rate (%)
100
80
60
40
20
0
Control1
2h_ RT1306
2h_ wtRT
2h_ SSIII
Stop Rate (%)
****
****
****
Deletion Rate (%)
**
****
Insertion Rate (%)

**c**

| Lib_ID | Description |
|---|---|
| lib1 | 20m_RT1306 |
| lib4 | 2h_RT1306 |

**N = 1088 non-Ψ sites**

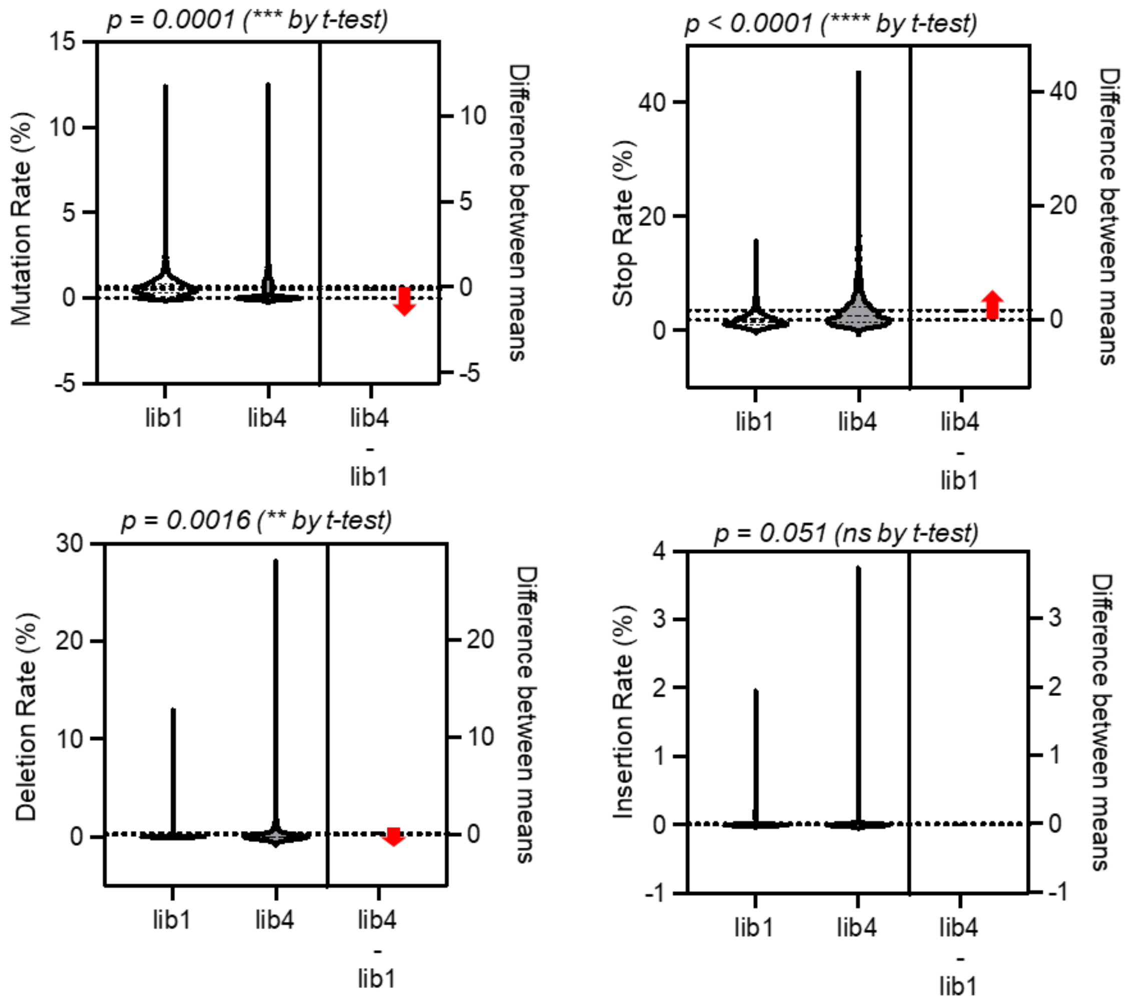


**Figure S7.** RT signature profiling based on reported Ψs and Us in the human rRNAs. a) Deletion rates of RT1306 towards CMC treated RNA, showing no obvious insertions b) RT signatures on unmodified Us by 3 different RTases in rRNAs via Amplicon sequencing libraries. Increased background noise with statistical significance is labeled. c) RT signatures on unmodified Us by RT1306 in rRNAs with 20-min and 2-hour CMC treatments. *P* values of figures were calculated by two-sided Student's t-test. Statistical significance: ns. = not significant, **P* < 0.05, ***P* < 0.01, ****P* < 0.001, *****P* < 0.0001.

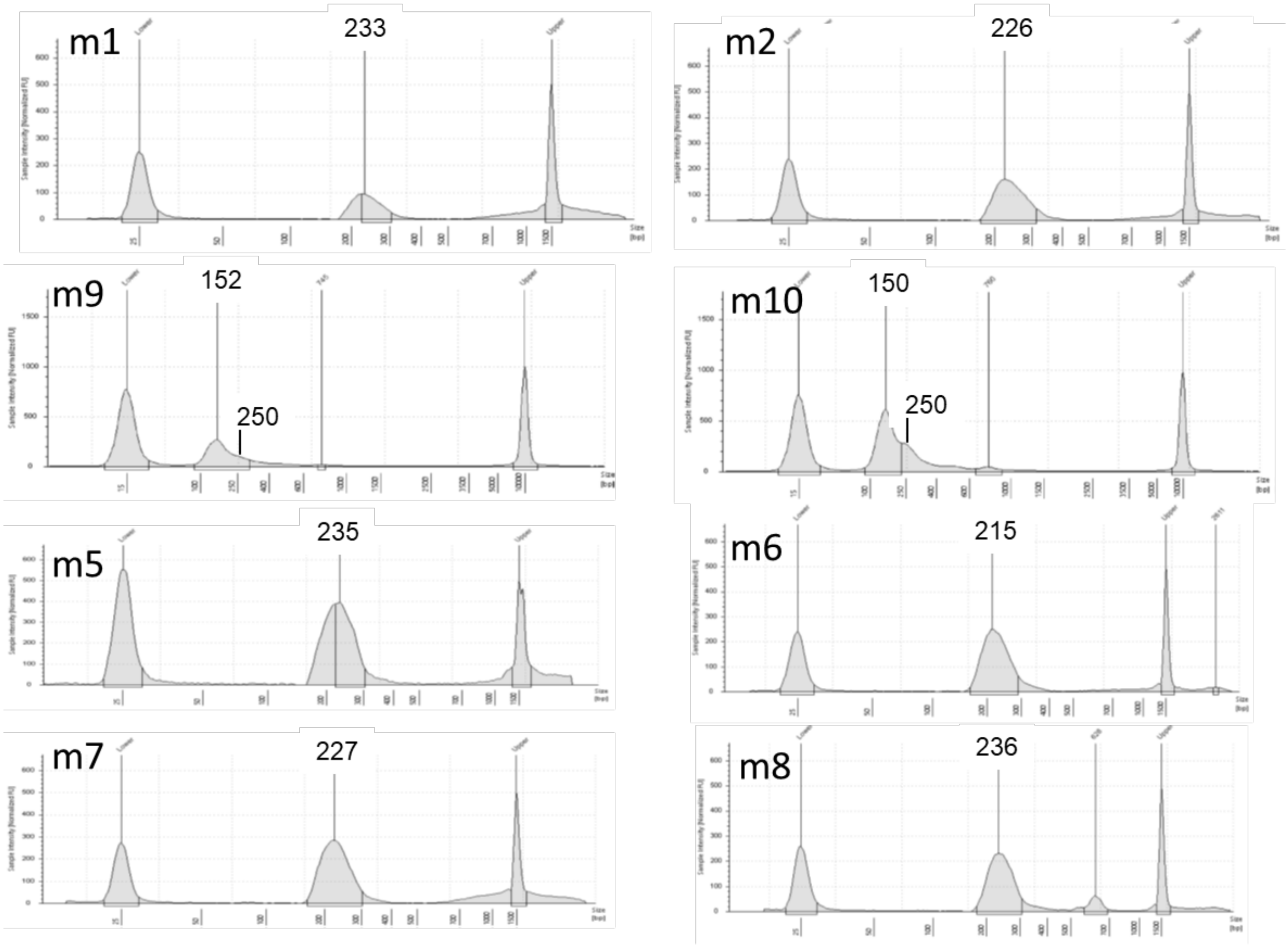


**Figure S8.** Mut-Ψ-seq libraries size analysis by Agilent TapeStation system.

a

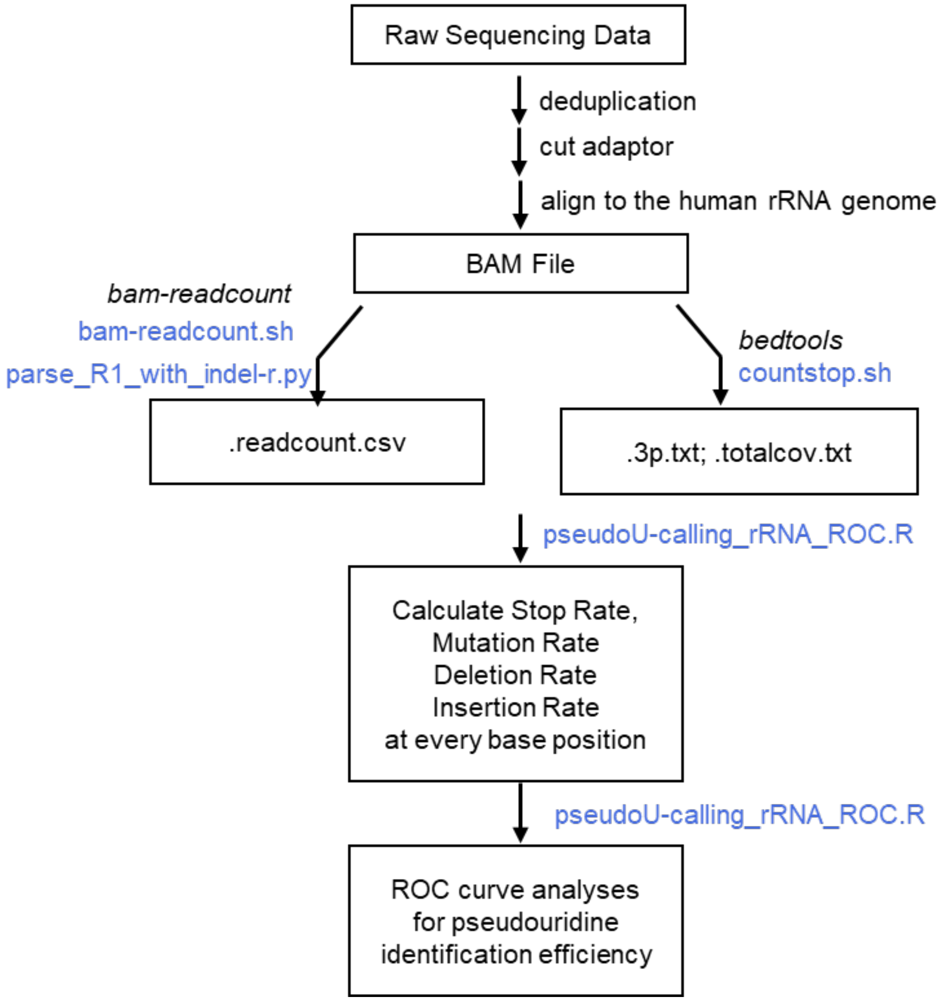
Raw Sequencing Data
deduplication
cut adaptor
align to the human rRNA genome
BAM File
bam-readcount
bam-readcount.sh
parse_R1_with_indel-r.py
bedtools
countstop.sh
.readcount.csv
.3p.txt; .totalcov.txt
pseudoU-calling_rRNA_ROC.R
Calculate Stop Rate,
Mutation Rate
Deletion Rate
Insertion Rate
at every base position
pseudoU-calling_rRNA_ROC.R
ROC curve analyses
for pseudouridine
identification efficiency

**b**

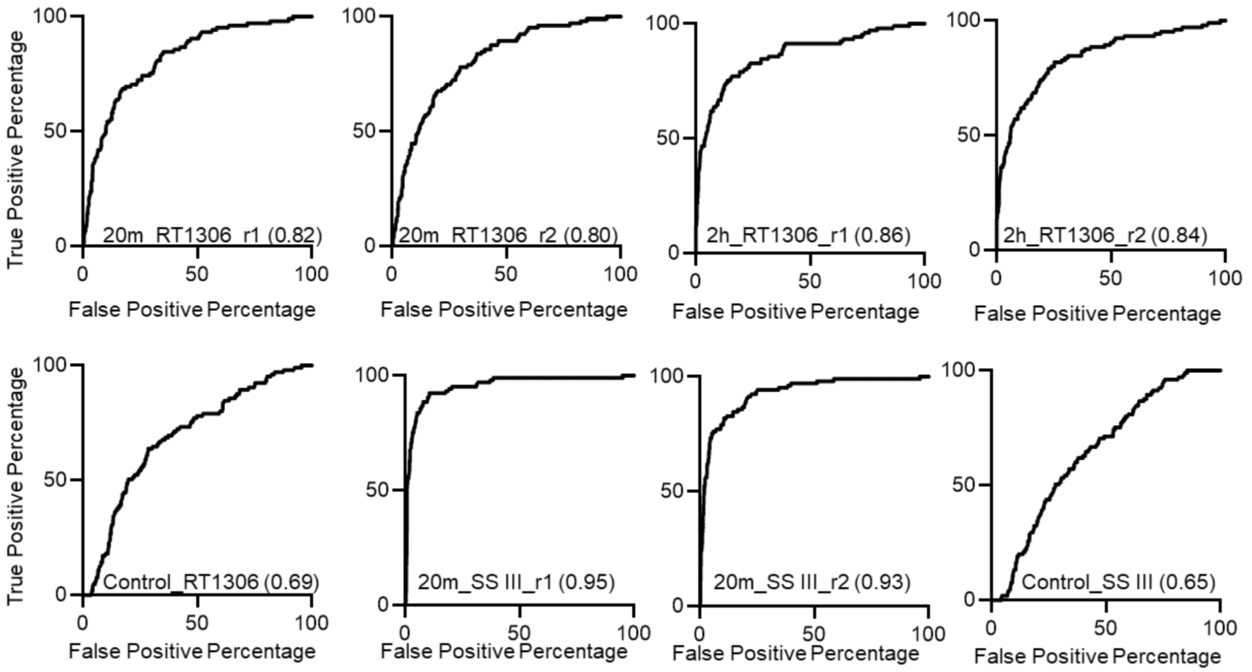
True Positive Percentage
False Positive Percentage
20m_RT1306_r1 (0.82)
20m_RT1306_r2 (0.80)
2h_RT1306_r1 (0.86)
2h_RT1306_r2 (0.84)
Control_RT1306 (0.69)
20m_SS III_r1 (0.95)
20m_SS III_r2 (0.93)
Control_SS III (0.65)

**c**

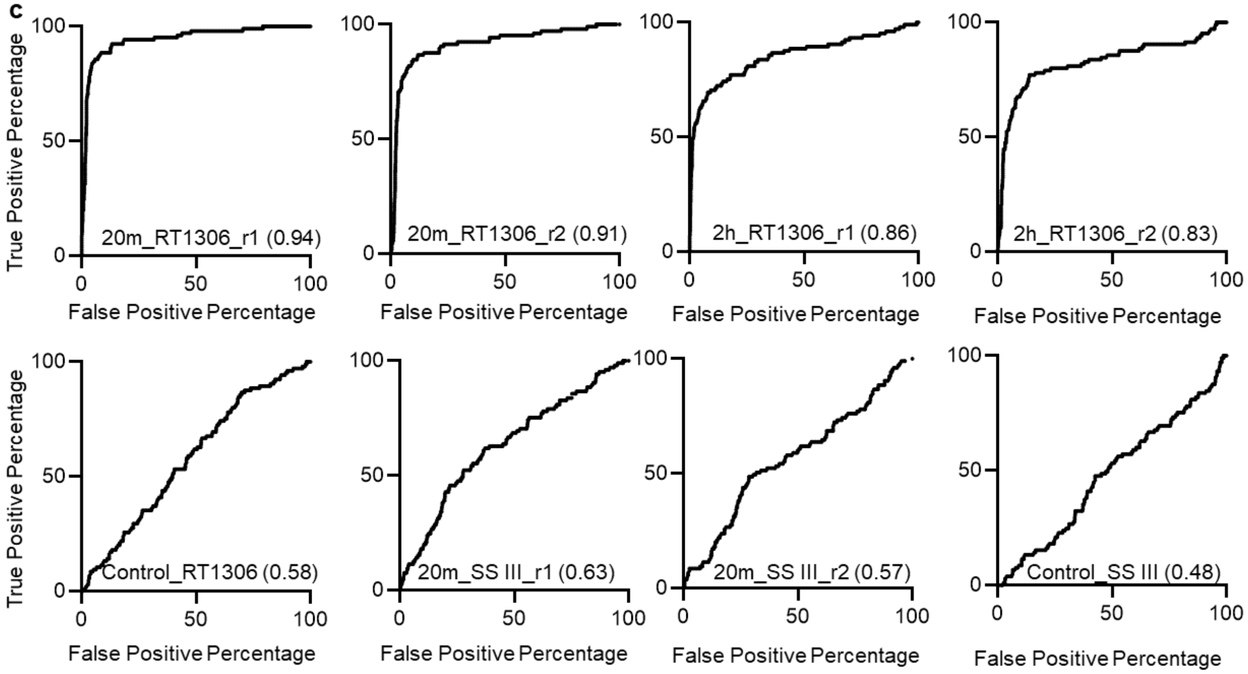
True Positive Percentage
False Positive Percentage
20m_RT1306_r1 (0.94)
20m_RT1306_r2 (0.91)
2h_RT1306_r1 (0.86)
2h_RT1306_r2 (0.83)
Control_RT1306 (0.58)
20m_SS III_r1 (0.63)
20m_SS III_r2 (0.57)
Control_SS III (0.48)

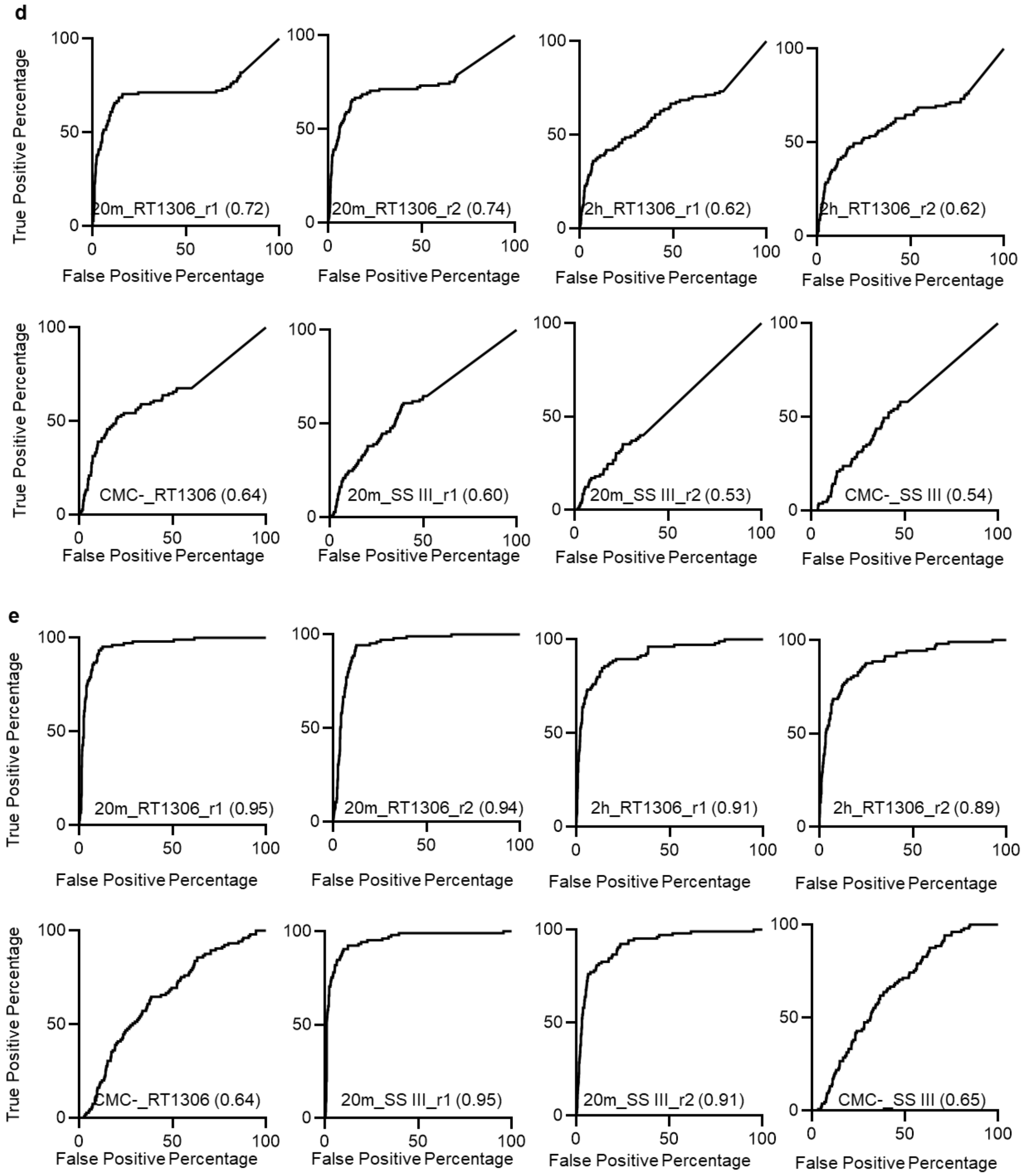


**Figure S9.** ROC analyses of RT signatures for Ψ identification in rRNAs with AUC indicated in the parathesis. a) Data processing pipeline for "piloting" Ψ-seq libraries. b) ROC analyses based on RT stops. c) ROC analyses based on RT mutations. d) ROC analyses based on RT deletions. e) ROC analyses based on combined RT signatures.

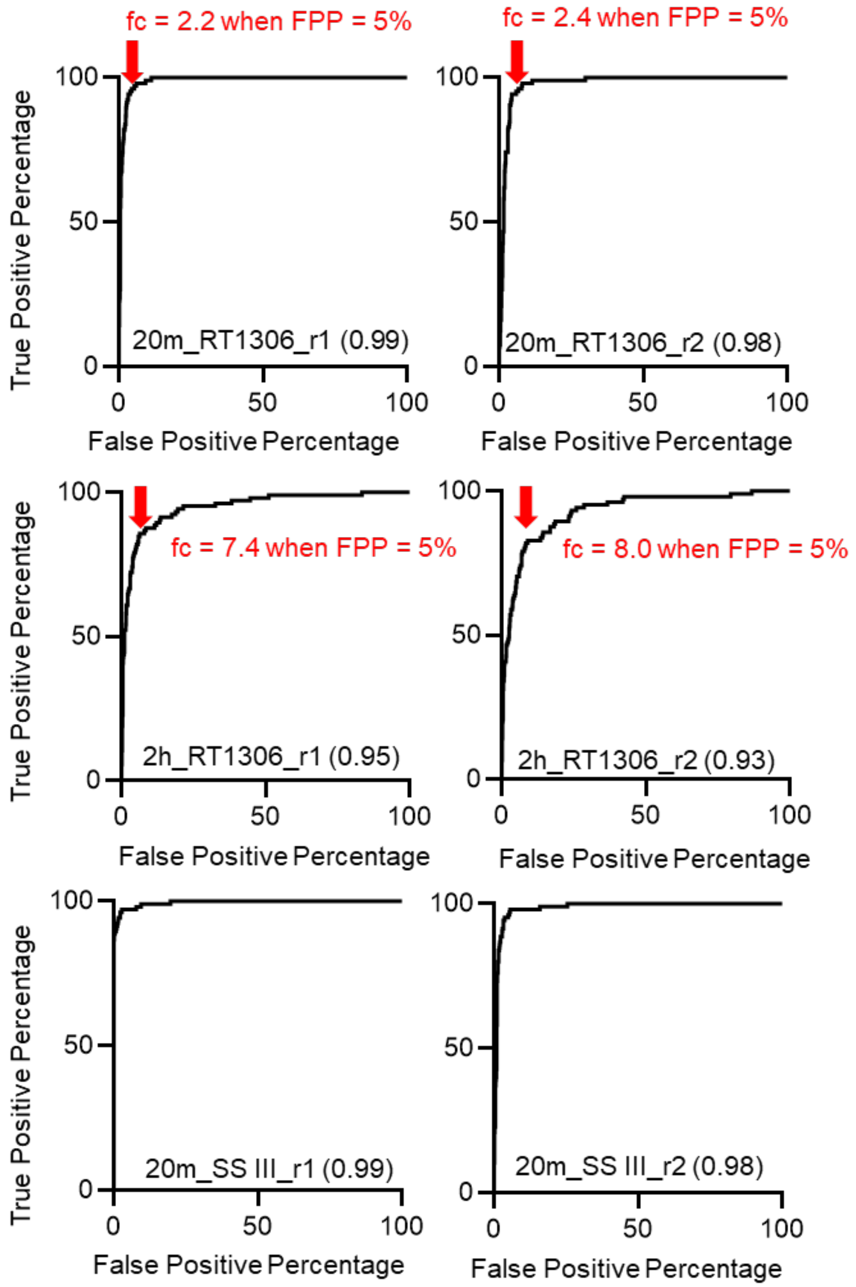


**Figure S10.** ROC analyses of the fold-change of the combined RT signatures for Ψ identification in rRNAs with AUC indicated in the parathesis. Fold changes (FC) at 5% false positive percentage (5% FPP) used for defining the cut-off threshold for Ψ identification are highlighted by red arrows with values indicated.

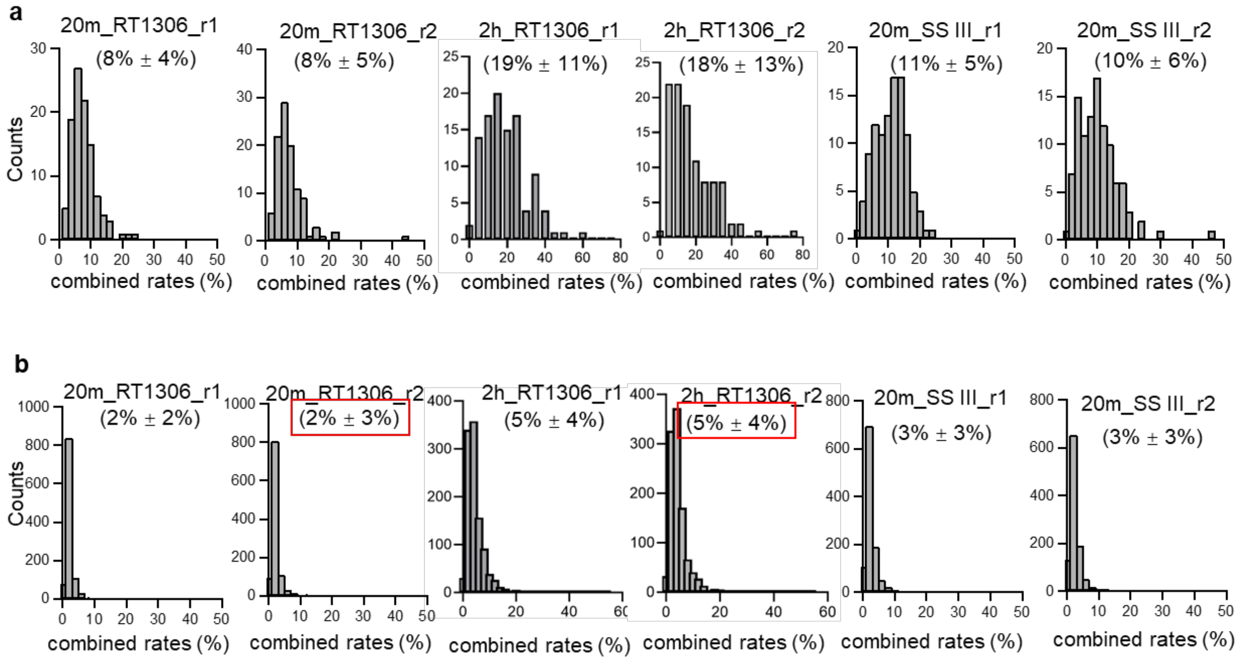


**Figure S11.** Histogram of the distribution of combined rates of a) Ψs in rRNA b) Us in rRNA in different sequencing libraries. Background levels of combined rates used for defining the cut-off threshold for Ψ identification are highlighted by red boxes: 5% for 20min-CMC treated libraries and 9% for 2hr-treated libraries.

a

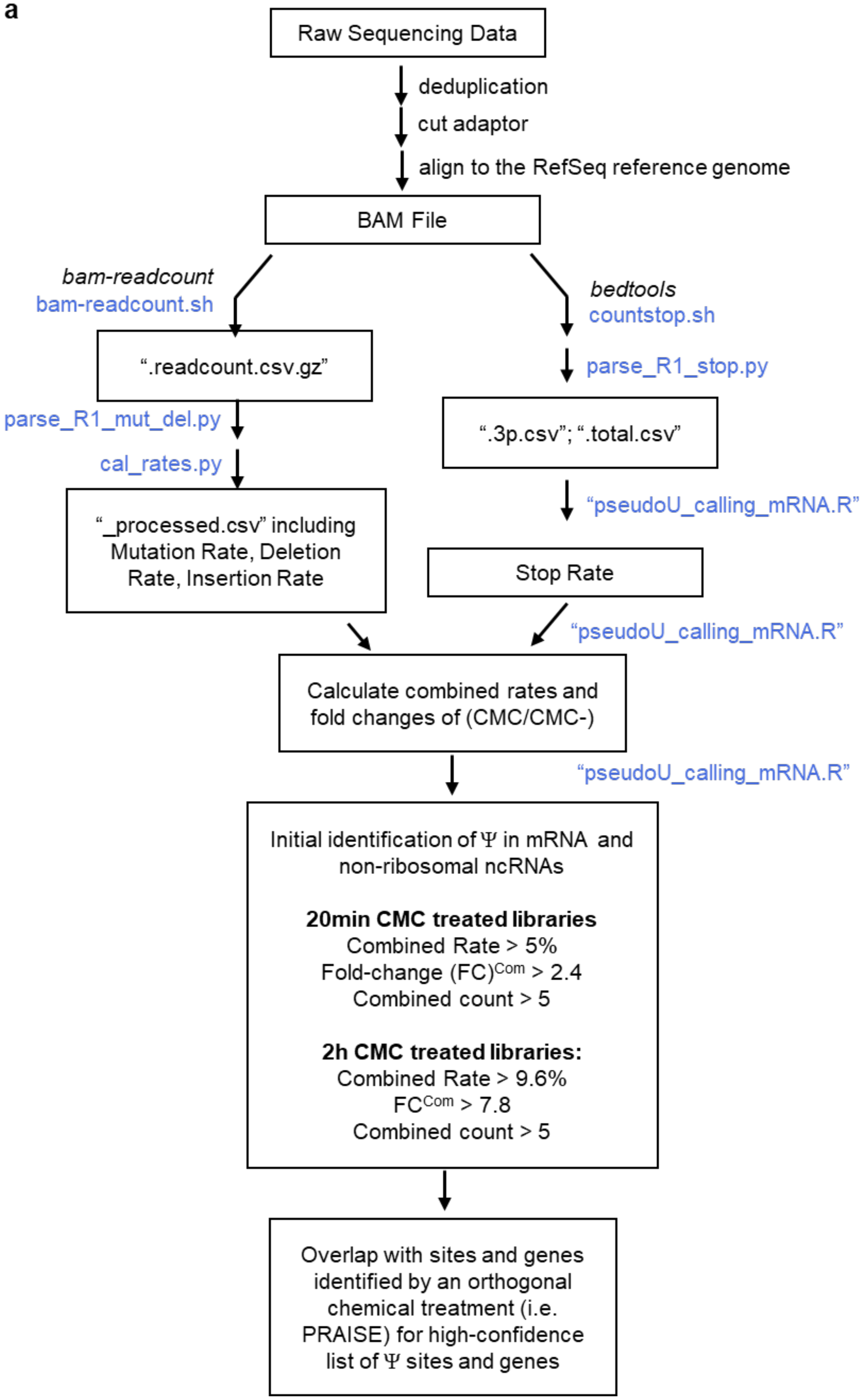
Raw Sequencing Data
deduplication
cut adaptor
align to the RefSeq reference genome
BAM File
bam-readcount
bam-readcount.sh
bedtools
countstop.sh
".readcount.csv.gz"
parse_R1_stop.py
parse_R1_mut_del.py
".3p.csv"; ".total.csv"
cal_rates.py
"_processed.csv" including Mutation Rate, Deletion Rate, Insertion Rate
"pseudoU_calling_mRNA.R"
Stop Rate
"pseudoU_calling_mRNA.R"
Calculate combined rates and fold changes of (CMC/CMC-)
"pseudoU_calling_mRNA.R"
Initial identification of Ψ in mRNA and non-ribosomal ncRNAs
20min CMC treated libraries
Combined Rate > 5%
Fold-change (FC)Com > 2.4
Combined count > 5
2h CMC treated libraries:
Combined Rate > 9.6%
FCCom > 7.8
Combined count > 5
Overlap with sites and genes identified by an orthogonal chemical treatment (i.e. PRAISE) for high-confidence list of Ψ sites and genes

**b**

| Lib_ID | Description | Treatment | RTase | Alignment Rate (%) |
|---|---|---|---|---|
| m1 | "20m_RT1306_r1" | CMC 20min; $OH^-$ | RT-1306 | 83.81 |
| m2 | "20m_RT1306_r2" | CMC 20min; $OH^-$ | RT-1306 | 84.13 |
| m9 | "2h_RT1306_r1" | CMC 2hr; $OH^-$ | RT-1306 | 61.43 |
| m10 | "2h_RT1306_r2" | CMC 2hr; $OH^-$ | RT-1306 | 42.23 |
| m5 | "Control_RT1306" | $OH^-$ | RT-1306 | 81.95 |
| m6 | "20m_SS III_r1" | CMC 20min; $OH^-$ | SS III | 87.89 |
| m7 | "20m_SS III_r2" | CMC 20min; $OH^-$ | SS III | 87.83 |
| m8 | "Control_SS III" | $OH^-$ | SS III | 89.95 |

**Figure S12.** a) Data processing pipeline for Mut-Ψ-seq. b) Alignment rate for the Mut-Ψ-seq libraries to the Refseq reference genome.

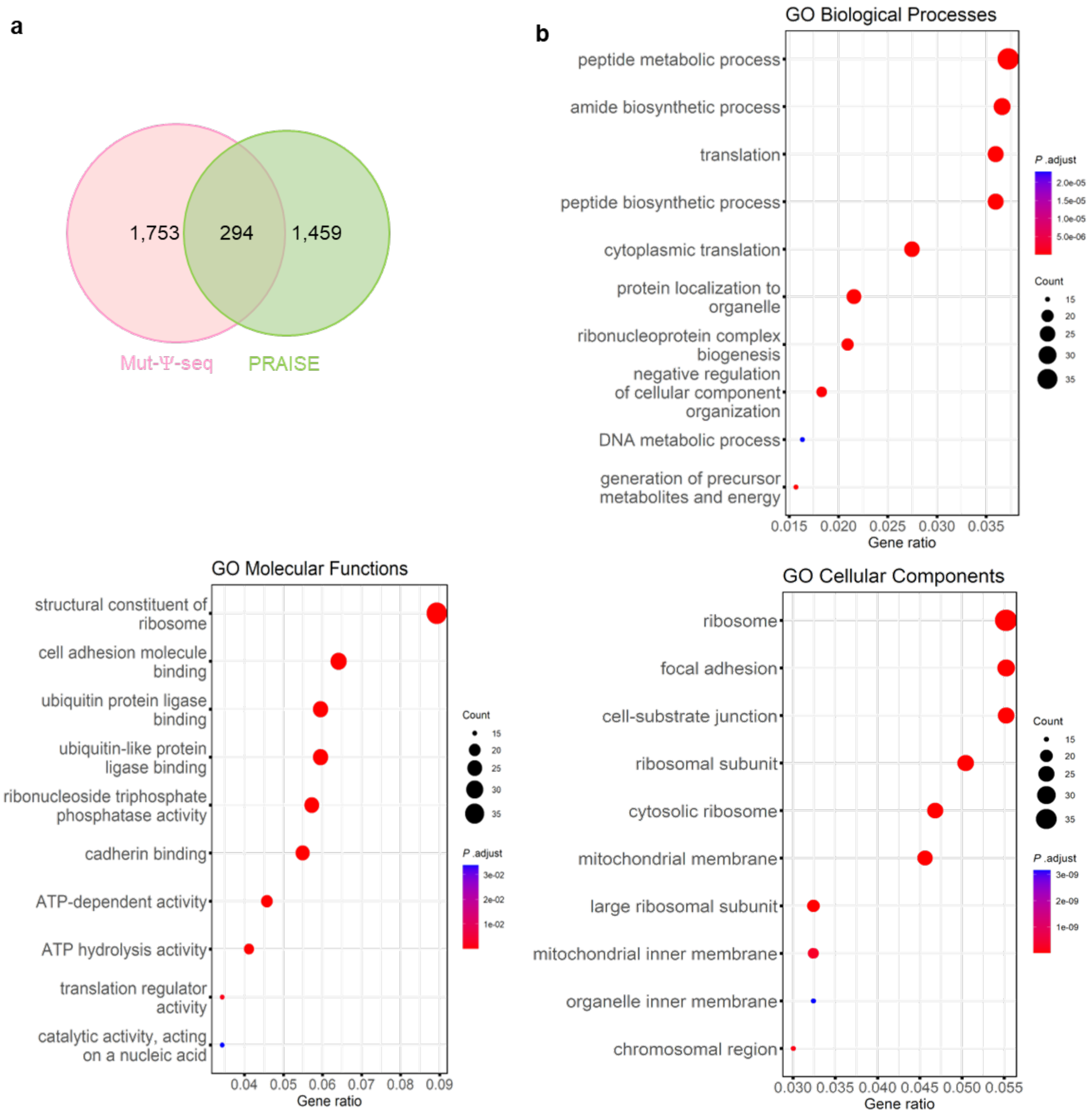


**Figure S13.** Comparison of mRNA Ψ sites identified by Mut-Ψ-seq with PRAISE. a) Venn diagram illustrating the overlap of mRNA Ψ sites between Mut-Ψ-seq (gene level) and PRAISE list. b) Gene ontology enrichment analysis (biological process, molecular functions, and cellular components) for 294 overlapped genes by orthogonal methods.

a

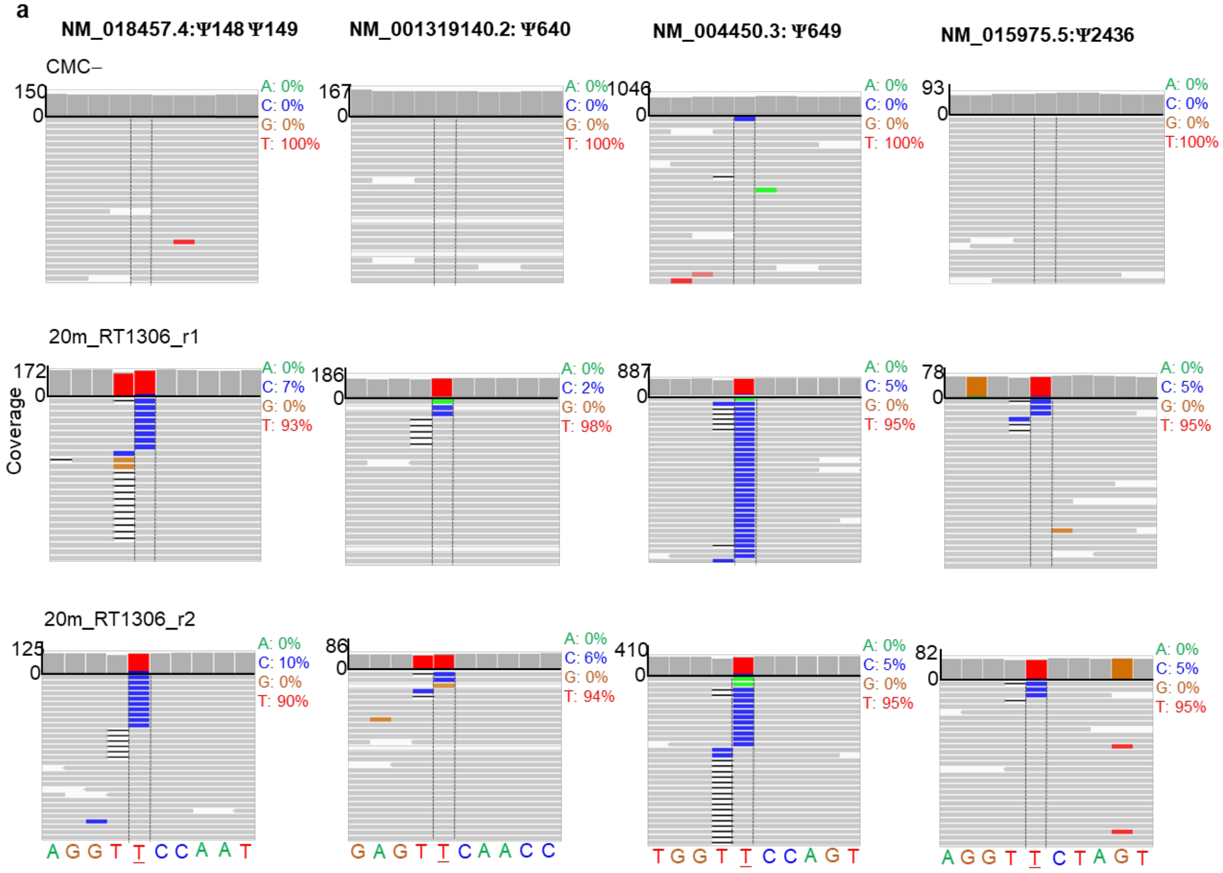
NM_018457.4:Ψ148 Ψ149
NM_001319140.2: Ψ640
NM_004450.3: Ψ649
NM_015975.5:Ψ2436
CMC–
150
0
A: 0%
C: 0%
G: 0%
T: 100%
167
0
A: 0%
C: 0%
G: 0%
T: 100%
1046
0
A: 0%
C: 0%
G: 0%
T: 100%
93
0
A: 0%
C: 0%
G: 0%
T:100%
20m_RT1306_r1
Coverage
172
0
A: 0%
C: 7%
G: 0%
T: 93%
186
0
A: 0%
C: 2%
G: 0%
T: 98%
887
0
A: 0%
C: 5%
G: 0%
T: 95%
78
0
A: 0%
C: 5%
G: 0%
T: 95%
20m_RT1306_r2
125
0
A: 0%
C: 10%
G: 0%
T: 90%
86
0
A: 0%
C: 6%
G: 0%
T: 94%
410
0
A: 0%
C: 5%
G: 0%
T: 95%
82
0
A: 0%
C: 5%
G: 0%
T: 95%
A G G T T C C A A T
G A G T T C A A C C
T G G T T C C A G T
A G G T T C T A G T

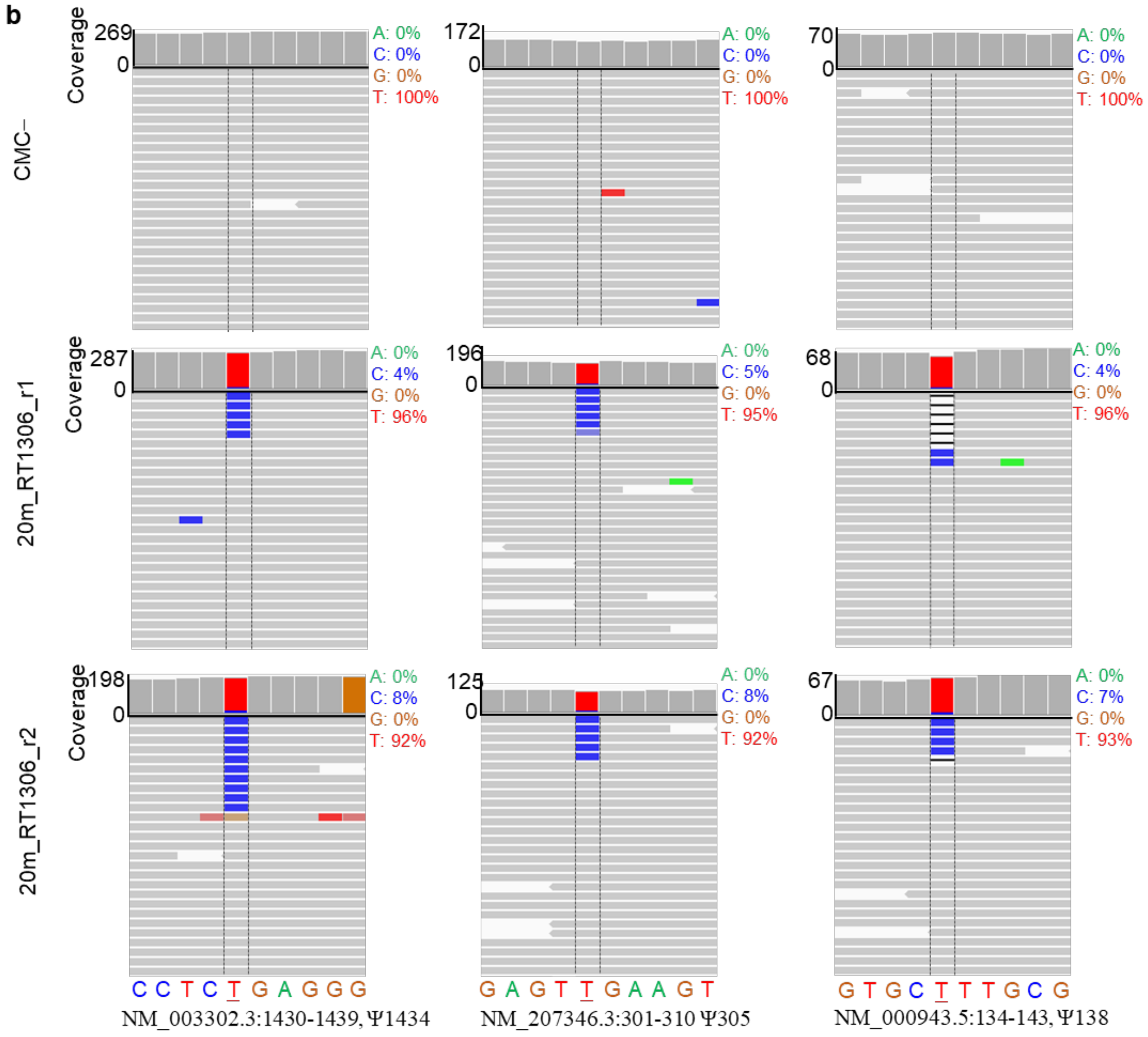


**Figure S14.** Promoted mutation signatures against CMC-Ψ in mRNAs by RT-1306. a) IGV views of 4 sites with GUUC context, b) IGV views of 3 examples with U, UU and UUU contexts.

### 2. Description of Processing Scripts

**Raw sequencing data processing**

- **"clump.sh":** Remove PCR duplication upon deduplication by "BBMap" tools "clumpify.sh" as the library carries the UMI sequence.

- **"cutadapt.sh":** remove Illumina adaptor sequences in the R1 reads.

**rRNA alignment and ROC curve analysis for identifying reported pseudouridine sites in rRNAs**

- **"bowtie2_r.sh"**: Align processed R1 reads to the human rRNA genes with Bowtie2. Output the resulting aligned reads as sorted bam files.

- **"bamreadcount.sh"**: count reads for all base types, insertion, and deletions at specific nucleotide positions. Outputs are in ".readcount.csv.gz".

- **"parse_R1_with_indel-r.py"**: Parse mutation, deletion, and insertion read counts from the bam-readcount outputs (".readcount.csv.gz" files) for all nucleotide positions. Outputs are in ".readcount.csv".

- **"countstop_r.sh":** count the reads that stop at each base position (output as ".3p.txt") and the total coverage at each base position (output as ".totalcov.txt") in the reference genome using "bedtools genomecov".

- **"pseudoU-calling_rRNA_ROC.R"**: Calculate the stop rate, mutation rate, deletion rate and insertion rate, then use the rRNA-Umod-annotations.txt as a reference to perform ROC curve analysis for any RT signatures: mutation rate, stop rate, deletion rate, combined rate, or fold-changes of combined rates.

**pseudouridine identification in mRNA and non-ribosomal ncRNAs**

- **"bowtie2_mRNA.sh":** Align processed R1 reads to the hg38-refseq reference genome with Bowtie2. Output the resulting aligned reads as sorted bam files.

- **"bam-readcount-m.sh":** count reads for all base types, insertion, and deletions at specific nucleotide positions. Outputs are in ".readcount.csv.gz".

- **"parse_R1_mut_del.py"**: Parse mutation, deletion, and insertion read counts from the bam-readcount outputs (".readcount.csv.gz" files) for nucleotide positions where the total read coverage is greater than 10 and the reference base is "T" or "t" in the reference genome. Outputs are in ".readcount.csv".

- **"cal_rates.py":** calculate mutation rate, deletion rate, and insertion rate at single base position from the ".readcount.csv" files. Output are as "_processed.csv"

- **"countstop_m.sh"**: count the reads that stop at each base position (output as ".3p.txt.gz") and the total coverage at each base position (output as ".totalcov.txt.gz") in the reference genome using "bedtools genomecov".

- **"parse_R1_stop.py":** parse out the stop counts and total coverage counts for nucleotide positions where the total read coverage is greater than 10 and the reference base is "T" or "t" in the reference genome, which resulted in output as ".3p.csv" and ".total.csv", respectively.

- **"pseudoU_calling_mRNA.R":** calculate the stop rate and the combined rates (stop + mutation + deletion rates) and call psedouridine sites in mRNAs and non-ribosomal ncRNAs by the combined rates of the CMC-treated libraries and the fold-change of combined rates of the CMC-treated versus nonCMC control libraries.

### 3. Description of the Supplementary Tables

**Table S1.** RNA and DNA oligonucleotides used in this research.
**Table S2.** Initial pseudouridine sites in mRNA and lncRNA called by combined rates fold change.
**Table S3.** The Gene ID of the overlapped genes by Mut-Ψ-seq and PRAISE.
